# Sleep resolves competition between explicit exemplar memory and implicit memory generalization

**DOI:** 10.1101/2025.02.21.639581

**Authors:** Katja Kleespies, Philipp C. Paulus, Hao Zhu, Florian Pargent, Marie Jakob, Jana Werle, Michael Czisch, Joschka Boedecker, Steffen Gais, Monika Schönauer

## Abstract

Sleep supports stabilization of explicit, declarative memory and benefits implicit statistical learning. In addition, sleep may change the quality of memory representations. Explicit and implicit learning systems, usually linked to hippocampus and striatum, can compete during learning, but whether they continue to interact during offline periods remains unclear. Here, we investigate for feedback-driven classification learning, whether sleep integrates explicit exemplar memory and implicit information-integration. The negative relationship between implicit and explicit memory components was resolved over sleep, but not wakefulness. Additionally, sleep benefitted performance on a categorization task that allows the use of both explicit and implicit strategies, and participants who slept showed superior performance in generalizing their knowledge to unseen exemplars. A reinforcement learning model relates this to better transfer of the learned exemplar value representation after sleep. Together, this suggests that sleep allows flexible access to information learned by different routes to respond optimally to everyday life contingencies.

## Introduction

To distinguish between actions with high and low reward, we must learn from different sources of information and combine them flexibly to guide adaptive behavior in novel situations. For instance, during childhood, we develop food preferences through explicit instruction from caregivers. At the same time, we learn about the underlying factors that determine food value by integrating knowledge from our own experiences. This dual learning process enables us not only to assess the value of familiar foods, but also to generalize our preferences to new and unfamiliar ones. Similarly, in classification learning tasks conducted in laboratory settings, two distinct types of representations are formed. Detailed explicit episodic memory of individual items allows generalization to unseen items based on similarity of these new items with previously stored exemplars (*exemplar-based* strategy) ^1^. Implicit statistical learning leads to a non-declarative (i.e., non-verbalizable) model of common relations between item features and associated responses, shaping behavior without awareness and developing gradually over a large number of learning trials (strategies based on implicit incremental learning, including *information-integration*) ^2,3^. While explicit memory is generally assumed to rely on structures in the medial temporal lobe, including the hippocampus, in interaction with prefrontal and posterior cortical regions ^4,5^, information-integration tasks rely on the striatum ^6,7^. Both memory systems can interact during task training, while the importance of systems for a specific task can shift over time ^8–10^. The nature of this interaction may be mediated by other brain regions including the prefrontal cortex ^11,12^. However, it remains unclear, which factors shape the interplay and integration of knowledge across these different memory systems. Here, we investigate whether sleep modulates the interaction between representations formed in explicit and implicit memory systems.

Sleep has been linked to memory consolidation and synaptic plasticity ^13,14^. It supports stabilization of explicit, declarative memory ^15–18^ and benefits implicit, procedural memory ^19–23^. Sleep is therefore considered crucial for the offline evolution of newly encoded memory traces ^24^. Information is not only selectively strengthened ^25,26^ (but see also ^27^), sleep-dependent memory processing has also been suggested to alter the quality of the memory trace ^28^. Sleep is thought to help item integration and multi-item generalization, leading to a functionally different representation of memories ^29–33^.

However, although evidence for sleep-dependent memory stabilization and enhancement in both the explicit and implicit domain is ample ^14^, the qualitative representational changes induced by sleep are still less well understood ^13,24,34^. Data from artificial grammar learning ^35–39^, statistical learning ^40,41^ and transitive inference ^42–46^ suggest that sleep in fact helps to extract implicit, hidden relations between studied items, leading to offline rule extrapolation. Furthermore, it has been suggested that sleep may induce gist abstraction in declarative learning tasks ^47–49^ and explicit insight into problem solving tasks ^50–52^.

These qualitative changes have been observed separately in implicit and explicit memory. However, many everyday learning tasks engage both implicit and explicit learning simultaneously, and it has been shown that these can compete during task training ^8,53,54^. Memory consolidation including overnight sleep seems to moderate the relationship between explicit and implicit learning systems ^41,55,56^. It remains unclear whether implicit and explicit aspects of a classification learning task are integrated over a night of sleep compared to wakefulness, or if they persist as separate representations. Recent evidence from spatial navigation learning indicates that sleep may favor the formation of a unified spatial memory by integrating hippocampal and striatal spatial representations ^57^. This suggests that sleep could resolve the competition between explicit and implicit memory systems in classification learning to allow flexible access to different memory representations depending on task demands.

In this study, we directly compared the contribution of sleep to the evolution of both explicit and implicit memory acquired during classification learning. We investigated a form of implicit contingency learning in a typical feedback-driven information-integration classification task in which participants learned a complex decision rule. Two different memory traces were established during learning: explicit knowledge about individual exemplars and implicit memory relying on integration and generalization of information regarding the underlying regularities. After periods of sleep or wakefulness, we tested explicit exemplar knowledge and implicit information-integration proficiency separately. Additionally, we tested participants on a categorization task that can be solved using both implicit and explicit memory.

## Results

Participants were trained on a classification learning task in which they had to distinguish between winning and losing exemplars (see Fig. 1). Participants performed 625 consecutive learning trials. On each given trial, two stimuli were presented on a black screen and participants had to choose the exemplar with the higher rank, i.e., the exemplar winning the respective comparison. The rank of the stimuli was defined by a hidden rule reflecting a linear combination of values assigned to the different stimulus properties (shape, symbol, fill color) while frame color served as an irrelevant distractor. Importantly, during learning, the same 25 exemplars were always the higher-ranking ones (winner exemplars) and the remaining 25 exemplars were always the lower-ranking ones (loser exemplars). Thus, participants could solve the task either by *explicitly* remembering if single exemplars were winners or losers (*exemplar-based decision*), or by *implicitly* learning and generalizing the underlying hidden regularities that determined the exemplar’s rank (decision based on *information-integration learning*). Participants were tested on different aspects of memory (implicit memory, explicit memory, and categorization based on both implicit and explicit memory) in a separate retrieval session. In the night-sleep and day-wake groups, participants were tested after a 12-h interval, during which they had either a full night of sleep or a normal day of wakefulness while participants in the circadian control groups (morning and evening control groups) were tested immediately after learning in the morning or evening, respectively.

**Fig. 1.**
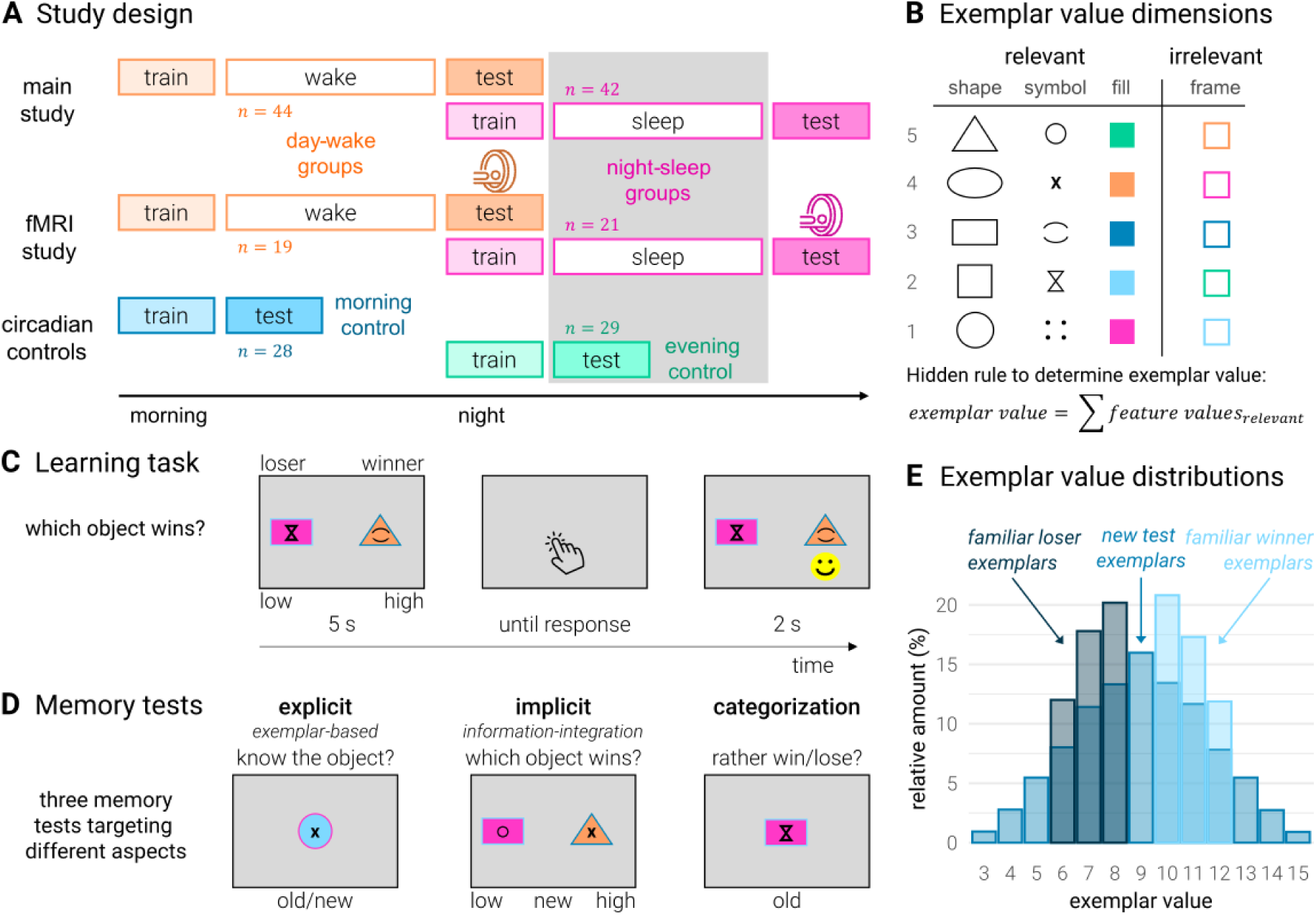
Study design and procedure. (A) Study design: *n* = 86 participants in the main study followed a 12-h day-wake (task training in the morning, testing in the evening) vs. 12-h night-sleep (task training at night, testing in the following day) design. For an additional *n* = 40 that followed the same schedule functional magnetic resonance imaging (fMRI) was acquired. To rule out circadian influences, *n* = 57 participants in the circadian control groups were either trained and tested in the morning (morning circadian control group) or evening (evening circadian control group). (B) Exemplar value dimensions: During the learning task, participants were presented exemplars that were assigned different values according to a hidden exemplar hierarchy unknown to the participants. Exemplar values were determined by adding the values (1-5) assigned to the properties (e.g., triangle, ellipse, rectangle, square, circle for the dimension ‘shape’) of the individual relevant dimensions (i.e., shape, symbol, fill color). The frame color served as a distractor. (C) Learning task: During task training, participants had to decide which of the two presented exemplars ranked higher according to the hidden value hierarchy, and to indicate the result via keypress with the left or right index finger. Correct decisions were reinforced with a happy face. Importantly, exemplars during training either always won (winner exemplars) or lost the comparisons (loser exemplars). Thus, participants could solve the task either by *explicitly* remembering for single exemplars if they were winners or losers (*exemplar-based decision*), or by *implicitly* learning and generalizing the underlying hidden regularities that determined the exemplar’s rank (decision based on *information-integration learning*). (D) Memory tests: Implicit information-integration memory for the hidden regularities was tested by pairing two new exemplars that had not been presented during training. Explicit exemplar memory was tested in a recognition task by presenting the 50 exemplars that had appeared during training and 50 novel exemplars. Exemplar categorization was assessed by asking participants to judge for individual, previously presented exemplars whether these exemplars would rather win or lose in comparison to other exemplars. (E) Exemplar value distributions: Winning exemplars during training ranked between 10 and 12, losing exemplars between 6 and 8. New exemplars presented during the implicit memory test spanned the whole possible exemplar sum point value range between 3 and 15.

### Participants learned to distinguish winning from losing exemplars

All participants successfully acquired knowledge about winning and losing exemplars during classification task training (Fig. 2A; Supplementary Tables S2A; S3A). As a result, they performed with high accuracy during the second half of task training (overall performance: 92.02 % ± 0.44 %; *t*(182) = 95.41, *p* < .001, *d* = 7.05 [one-sample *t*-test against chance performance]; night-sleep group: 91.54 % ± 0.67 %; day-wake group: 92.27 % ± 0.80 %; morning control group: 91.98 % ± 1.38 %; evening control group: 92.53 % ± 0.93 %). Importantly, performance during the second half of training did neither differ between participants that were tested immediately after learning (morning ≠ evening circadian control group: *t*(55) = −0.33, *p* = .740, *d* = −0.09), nor between participants that received the delayed tests (night-sleep ≠ day-wake group: *t*(124) = −0.70, *p* = .487, *d* = −0.12), nor between participants that learned the classification task during the same time of day (night-sleep ≠ evening circadian control group: *t*(90) = −0.84, *p* = .403, *d* = −0.19; day-wake ≠ morning circadian control group: *t*(89) = 0.19, *p* = .847, *d* = 0.04). Because participants could apply different learning strategies during training, we fitted a reinforcement learning (RL) model on behavioral choices during the learning task to predict their behavior during the later memory tests assessing the use of explicit and implicit memory representations (Fig. 2B; Supplementary Tables S4A; S5A; see ‘Methods’). To probe validity of the applied modelling approach, we performed a recovery analysis where we predicted participants’ choices during learning from the trained model parameters. Overall, the model was able to predict participants’ behavior with high accuracy (overall performance: 85.44 % ± 0.45 %; *t*(182) = 78.44, *p* < .001, *d* = 5.80 [one-sample *t*-test against chance performance]; night-sleep group: 84.84 % ± 0.70 %; day-wake group: 85.77 % ± 0.83 %; morning control group: 85.23 % ± 1.46 %; evening control group: 86.23 % ± 0.83 %; Fig. 2C). Importantly, these prediction accuracies again did not differ either between participants who were tested immediately after learning (morning ≠ evening circadian control group: *t*(55) = −0.60, *p* = .549, *d* = −0.16), or between participants who received the delayed tests (night-sleep ≠ day-wake group: *t*(124) = −0.86, *p* = .393, *d* = −0.15), or between participants who learned the classification task during the same time of day (night-sleep ≠ evening circadian control group: *t*(90) = −1.18, *p* = .240, *d* = −0.27; day-wake vs. morning circadian control group: *t*(89) = 0.34, *p* = .733, *d* = 0.08). Across all learning trials, participants’ RL weights for the features relevant for classification exhibited a selective, incremental increase which was not observed for the irrelevant feature (Fig. 2D). This resulted in significantly different weights for the relevant features compared to the irrelevant feature at the end of learning (shape: 0.48 ± 0.01; *t*(182) = 44.16, *p* < .001; symbol: 0.48 ± 0.01; *t*(182) = 40.34, *p* < .001; fill: 0.48 ± 0.01; *t*(182) = 41.41, *p* < .001; frame (irrelevant): 0.00 ± 0.00; Fig. 2E; Supplementary Tables S12A, S12B). Together, this demonstrates that the participants became proficient in correct classification and that our RL model is suitable to predict behavioral choices during classification.

**Fig. 2.**
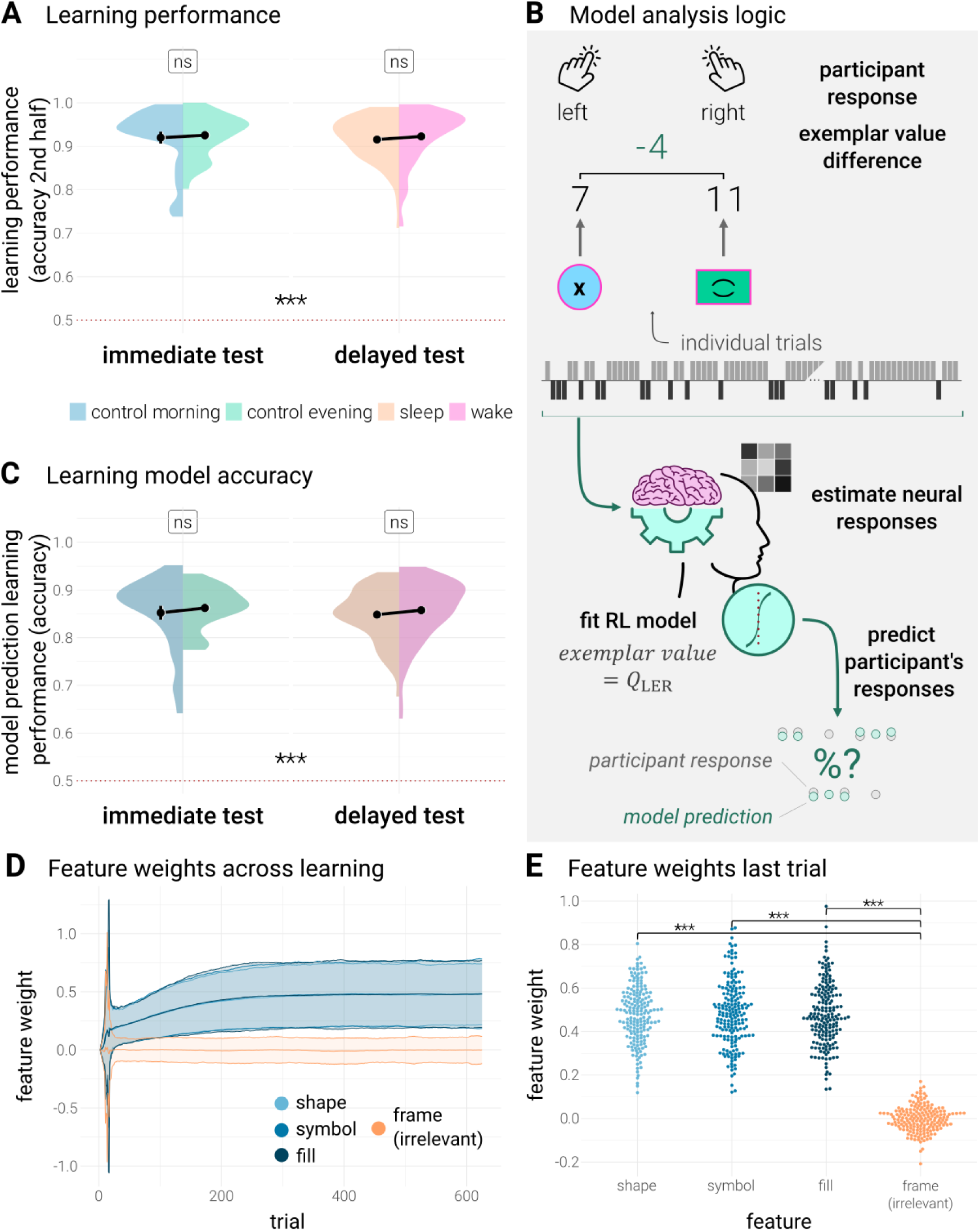
Task learning performance and reinforcement learning (RL) model. (A) Learning performance: Participants successfully acquired knowledge about winning and losing exemplars during classification task learning, indicated by overall above chance accuracy in the second half of learning. Performance did not differ between groups. (B) Model analysis logic: Participants’ behavior during task training and memory tests was modeled using a RL algorithm. The model was fitted to participants’ behavioral choices during task training and provided the learned exemplar value representation (*Q*-values) for each participant individually. Based on this model, participants’ responses during the memory tests were predicted using different policies for the explicit, the implicit, and the categorization memory task to assess whether participants were able to successfully apply the learned exemplar and corresponding value representations to solve the respective memory tasks. Additionally, the exemplar value representation was related to neural responses measured with functional magnetic resonance imaging (fMRI). (C) Model accuracy in the learning task was consistently high, confirming the validity of our model approach. No differences between groups were observed. (A, C) Data are *M* ± *SEM*. Red dotted lines indicate chance performance (with asterisks above line indicating significant difference from chance performance), a value of 1 indicates 100 % accuracy. (D) Feature weights across learning: Participants exhibit a selective, incremental increase for features relevant for classification (*M* ± 2 *SD*). (E) Feature weights last learning trial: Feature weights across participants at the end of learning are higher for all relevant compared to the irrelevant feature. ***: *p* < .001; ns: *p* > .1.

### Participants successfully acquired explicit and implicit memory representations during classification task training

The main objective of this study was to test if sleep resolves the competition between explicit exemplar memory and implicit information-integration proficiency. For this, participants were tested separately on their explicit and their implicit memory in the retrieval session, as well as on a categorization test that could be solved using both explicit and implicit memory strategies (see Fig. 1D). We first confirmed that participants were able to acquire both explicit exemplar and implicit information-integration rule representations during classification task training. Explicit exemplar memory was tested in a recognition task contrasting the exemplars that had appeared during learning with new ones. Implicit memory for generalization of the hidden regularities was tested by pairing two new exemplars. Participants had to decide which exemplar ranked higher. Known exemplar categorization was assessed by presenting exemplars studied during the learning task and asking participants whether an exemplar would rather win or lose in an exemplar comparison. In this task, best performance can be achieved by relying on a combination of explicit and implicit memory representations. Overall, participants performed well in all three tasks (overall explicit performance [*d*’]: 0.86 ± 0.03; *t*(182) = 25.35, *p* < .001, *d* = 1.87 [one-sample *t*-test against chance performance]; Supplementary Tables S2B, S3B; overall implicit performance [accuracy]: 65.28 % ± 0.54 %; *t*(182) = 28.43, *p* < .001, *d* = 2.10 [one-sample *t*-test against chance performance]; Supplementary Tables S2C, S3C; overall known exemplar categorization performance [accuracy]: 80.30 % ± 0.65 %; *t*(182) = 46.94, *p* < .001, *d* = 3.47 [one-sample *t*-test against chance performance]; Supplementary Tables S2D, S3D).

### Sleep resolves competition between explicit and implicit memory systems

We next analyzed whether there was a tradeoff between explicit and implicit memory performance (Supplementary Table S1C). When participants were tested immediately after learning (circadian control groups), the relationship between explicit and implicit aspects of their memory was competitive, as indicated by a significant negative relationship between explicit and implicit memory scores (morning control group: *b* = −0.09 ± 0.03, *t*(175) = −3.25, *p* = .001; evening control group: *b* = −0.09 ± 0.03, *t*(175) = −2.91, *p* = .004; Fig. 3A). However, this initially competitive relationship was selectively resolved after a 12 h-consolidation interval including sleep (night-sleep group: *b* = 0.01 ± 0.02, *t*(175) = 0.74, *p* = .460; Fig. 3B). Importantly, we did not observe this effect after a time interval of the same length of wakefulness (day-wake group: *b* = −0.08 ± 0.02, *t*(175) = −3.55, *p* < .001). Consequently, regression slopes differed significantly between night-sleep and day-wake groups, but not between circadian control groups (night-sleep ≠ day-wake: *b* = 0.09 ± 0.03, *t*(175) = 3.09, *p* = .002; morning control ≠ evening control: *b* = 0.00 ± 0.04, *t*(175) = −0.10, *p* = .923) while the difference between night-sleep and day-wake groups trended to be significantly larger than the difference between circadian control groups (night-sleep – day-wake ≠ control morning – control evening: *b* = 0.09 ± 0.05, *t*(175) = 1.88, *p* = .062). This demonstrates that an initially competitive relationship between explicit and implicit memory systems is selectively resolved by sleep. Note, that we did not only observe this sleep-dependent resolution of competition in the large study sample that combines data from behavioral and functional magnetic resonance imaging (fMRI) study protocols (Fig. 1A, see ‘Methods’), but also in the individual study samples, independently (Supplementary Tables S1A-C).

**Fig. 3.**
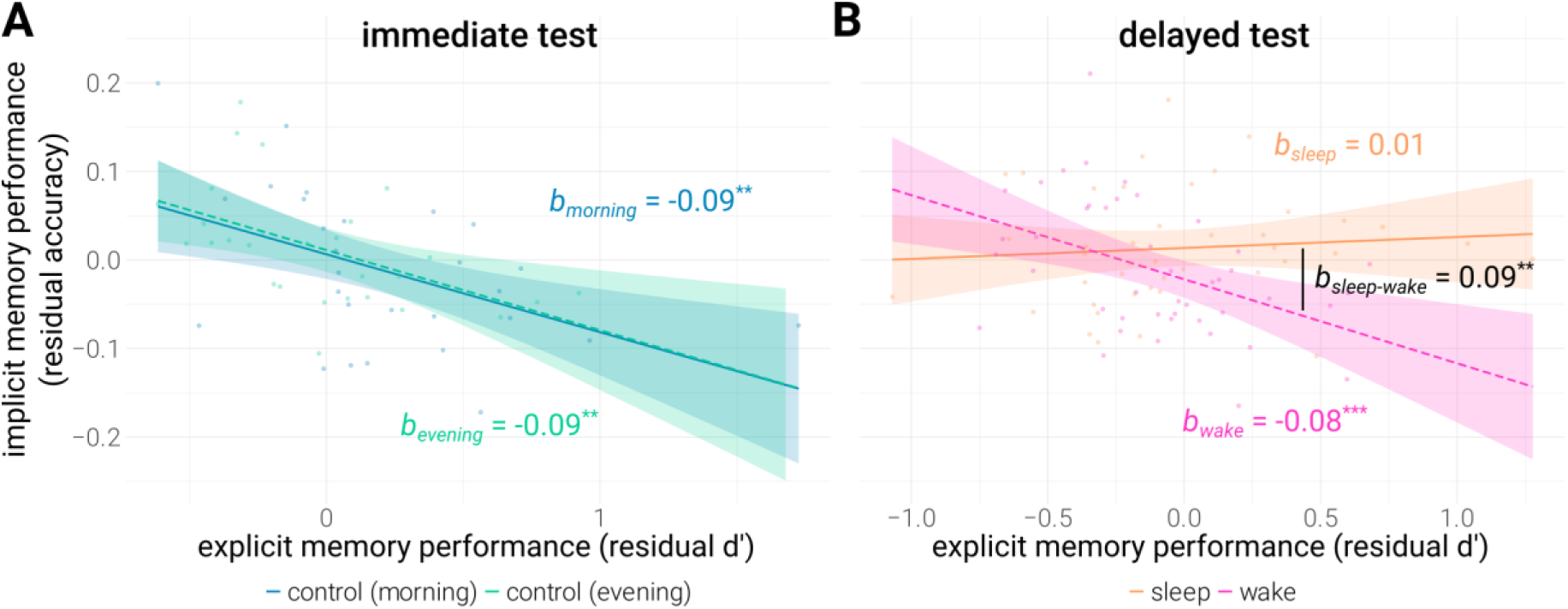
Relationship between explicit and implicit memory performance. Relationship between performance in explicit and implicit memory tests at (A) immediate tests (morning and evening circadian control groups) and (B) after a consolidation interval of 12 h (night-sleep and day-wake groups). Immediately after learning, the relationship between explicit and implicit aspects of their memory was competitive, i.e., negatively correlated, but was selectively resolved after a 12 h-consolidation interval including sleep and not wakefulness. (A, B) Regression line ± 95 %-confidence interval. Memory scores are displayed as residual scores after regressing out learning performance in the second half of task training. ***: *p* < .001; **: *p* < .01.

### Sleep benefits generalization of the exemplar value rule to unseen exemplars

Resolved competition between explicit and implicit memory systems after sleep might influence overall performance at test by allowing flexible access to the most effective decision strategy at the moment. Consequently, we assessed general group differences between explicit recognition, implicit generalization, and known exemplar categorization memory scores (Fig. 4). A night of sleep significantly improved implicit information-integration learning compared to a day of wakefulness, circadian effects were not observed (night-sleep [0.02 ± 0.01] ≠ day-wake group [−0.01 ± 0.01]: *t*(124) = 2.51, *p* = .014, *d* = 0.45; morning control [−0.01 ± 0.02] ≠ evening control group [0.01 ± 0.01]: *t*(55) = −1.20, *p* = .234, *d* = −0.32; Fig. 4B; Supplementary Tables S2C, S3C). Similarly, when explicit and implicit knowledge could be used in the same categorization task, we again found a positive effect of sleep over wakefulness while we did not find significant differences in performance between the morning and evening groups in this task (night-sleep [0.01 ± 0.01] ≠ day-wake group [−0.02 ± 0.01]: *t*(124) = 2.79, *p* = .006, *d* = 0.50; morning control [0.01 ± 0.01] ≠ evening control group [0.03 ± 0.01]: *t*(55) = −0.97, *p* = .336, *d* = −0.26; Fig. 4C; Supplementary Tables S2D, S3D). Explicit recognition of old exemplars was not affected by sleep and trended to be better in the morning control group (night-sleep [−0.03 ± 0.05] ≠ day-wake group [−0.11 ± 0.05]: *t*(124) = 1.14, *p* = .256, *d* = 0.20; morning control [0.26 ± 0.09] ≠ evening control group [0.04 ± 0.07]: *t*(55) = 1.90, *p* = .063, *d* = 0.50; Fig. 4A; Supplementary Tables S2B, S3B). Sleep thus increased implicit knowledge about the exemplar hierarchy rule and its generalization to new unseen exemplars as well as performance in the categorization task where this knowledge can be used jointly with explicit memory representations.

**Fig. 4.**
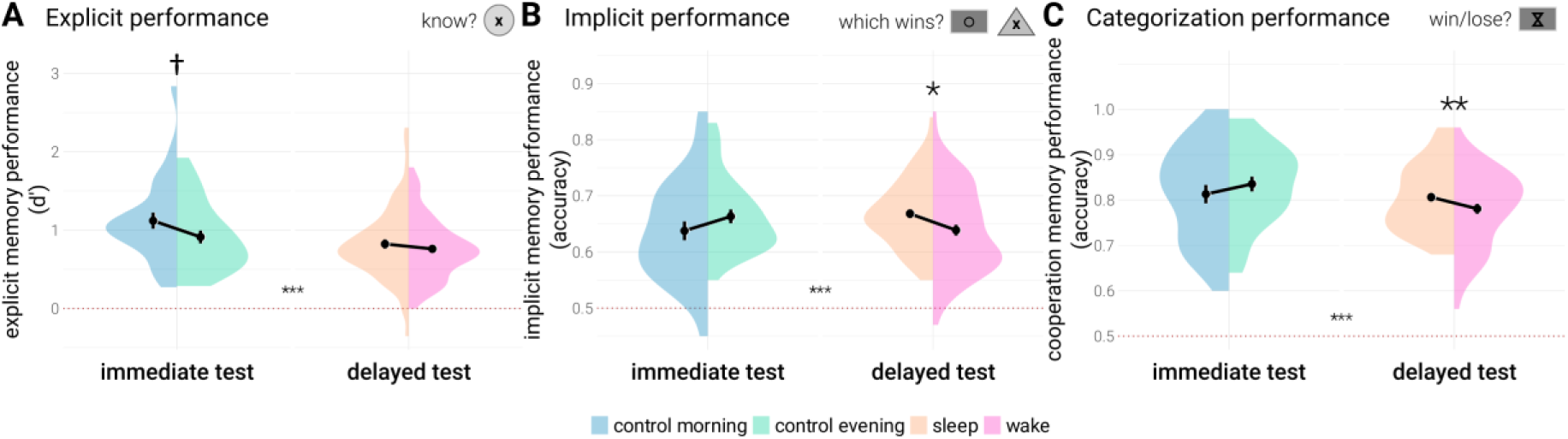
Behavioral performance in memory tests. Group comparisons of behavioral performance in the explicit, implicit, and categorization memory tests. (A) Explicit performance: Explicit recognition of old items was not affected by sleep, yet trended to be better in the morning. (B) Implicit performance: A night of sleep significantly improved implicit information-integration learning compared to a day of wakefulness while no circadian effects were observed. (C) Categorization performance: Similarly, when explicit and implicit knowledge could be used in the same test, we again found a positive effect of sleep over wakefulness while no circadian effects were observed. (A to C) Data are *M* ± *SEM*. To aid comprehension, uncorrected memory scores are displayed although statistical tests were performed on residual scores after regressing out learning performance in the second half of task training. Red dotted lines indicate chance performance (with asterisks above line indicating significant difference from chance performance). (B, C) A value of 1 indicates 100 % accuracy. ***: *p* < .001; **: *p* < .01; *: *p* < .05; †: *p* < .1.

### Sleep promotes transfer of the exemplar value representation acquired during task training

To better understand the mechanisms underlying the resolved competition between memory systems after sleep as well as better generalization of the information-integration rule, we again made use of the RL model fitted on participants’ learning task performance, to model choices in the memory tests (Fig. 5; see Fig. 2B, ‘Methods’). This allowed us to investigate the influence of sleep on participants’ memory representations by applying different decision policies to the learned exemplar value representation depending on the memory test used. In the explicit recognition memory test, we applied a policy for which the probability to rate an exemplar as ‘old’ increased with the number of exemplar features matching a specific exemplar from the learning exemplars. This policy was overall able to accurately predict participants’ performance on this task (overall explicit model accuracy: 69.82 % ± 0.64 %; *t*(182) = 109.95, *p* < .001, *d* = 8.13 [one-sample *t*-test against chance performance]) and yielded no differences between the night-sleep and day-wake group in accurately predicting participants’ choices in the explicit memory test (night-sleep [70.98 % ± 1.17 %] ≠ day-wake group [69.86 % ± 1.00 %]: *t*(124) = 0.73, *p* = .464, *d* = 0.13; Fig. 5A). Likewise, we did not observe circadian effects on the prediction of participants’ choices in the explicit memory test (control morning [69.71 % ± 1.86 %] ≠ control evening group [67.31 % ± 1.29 %]: *t*(55) = 1.07, *p* = .290, *d* = 0.28; Fig. 5A; Supplementary Tables S4B, S5B). Accordingly, sleep did not seem to influence transfer of explicit exemplar representations from task training to testing in this memory task. Importantly, we also tested whether sleep aids transfer of the acquired value representation from learning to the implicit memory generalization test. We applied a policy that recombined the individual feature value representations obtained from the RL model to predict participants’ behavioral choices in the implicit memory test (overall implicit model accuracy: 65.35 % ± 0.54 %; *t*(182) = 28.64, *p* < .001, *d* = 2.10 [one-sample *t*-test against chance performance]). Here, we observed better transfer of the acquired value representation to unknown exemplars in the night-sleep compared to the day-wake group while no circadian effects were observed (night-sleep [66.87 % ± 0.77 %] ≠ day-wake group [63.98 % ± 0.97 %]: *t*(124) = 2.33, *p* = .021, *d* = 0.42; control morning [64.00 % ± 1.69 %] ≠ control evening group [66.31 % ± 1.22 %]: *t*(55) = −1.12, *p* = .269, *d* = −0.30; Fig. 5B; Supplementary Tables S4C, S5C). Furthermore, predicting participants’ choices in the known exemplar categorization task using a value-based policy (overall known exemplar categorization model accuracy: 80.31 % ± 0.64 %; *t*(182) = 47.06, *p* < .001, *d* = 3.48 [one-sample *t*-test against chance performance]) showed a trend towards better transfer of the acquired value representation from learning to testing in the night-sleep compared to the day-wake group. Here, the policy applied to the model was more likely to indicate that an exemplar would rather win than lose in an exemplar comparison if the if the participants’ acquired exemplar value ranked high within all possible exemplars. Accordingly, prediction of participants’ choices in the known exemplar categorization task in the night-sleep group tended to be more accurate than in the day-wake group (night-sleep [80.60 % ± 0.94 %] ≠ day-wake group [78.06 % ± 1.07 %]: *t*(124) = 1.78, *p* = .077, *d* = 0.32; Fig. 5C). No circadian effects were observed (control morning [81.29 % ± 2.02 %] ≠ control evening group [83.59 % ± 1.62 %]: *t*(55) = −0.89, *p* = .377, *d* = −0.24; Fig. 5C; Supplementary Tables S4D, S5D). In summary, sleep allows for better transfer of the learned exemplar value representation to later memory tests. This better transfer might, in turn, facilitate rule generalization to new exemplars and resolve competition between memory systems.

**Fig. 5.**
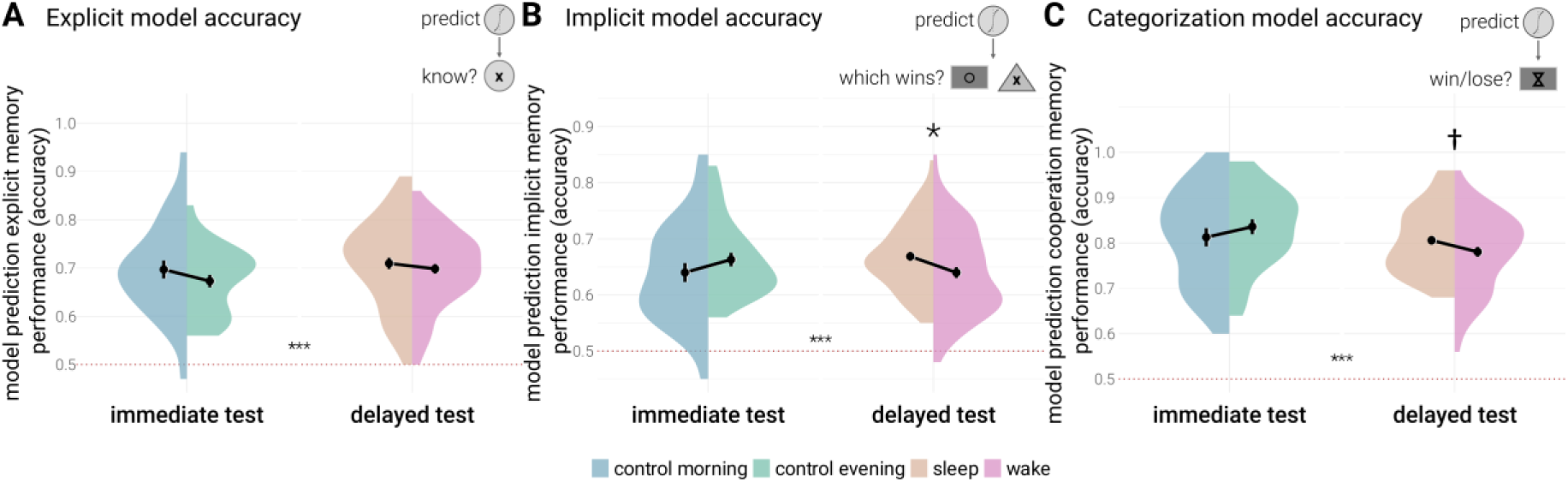
Reinforcement learning (RL) model prediction of memory tests. Group comparisons predicting participants’ behavior in the explicit, implicit, and cooperation memory tests employing different policies of the RL model. (A) Explicit model accuracy: No differences between groups in accurately predicting participants’ choices in the explicit memory test were observed. (B) Implicit model accuracy: Participants in the night-sleep group demonstrated better transfer of the acquired value representation from learning to testing compared to the day-wake group. This was reflected in better prediction of participants’ choices in the implicit memory test in the night-sleep compared to the day-wake group, while no circadian effects were observed. (C) Categorization model accuracy: Similarly, using a value-based policy, prediction of participants’ choices in the known exemplar categorization test in the night-sleep group trended to be more accurate than in the day-wake group while no circadian effects were observed. (A to C) Data are *M* ± *SEM*. Red dotted lines indicate chance performance (with asterisks above line indicating significant difference from chance performance), a value of 1 indicates 100 % accuracy. ***: *p* < .001; *: *p* < .05; †: *p* < .1.

Sleep is thought to support item integration and multi-item generalization, leading to a functionally different representation of memories ^29–33^. Accordingly, sleep may benefit the transfer of the learned exemplar value representation by supporting a combined, multi-dimensional use of the acquired feature values. Alternatively, sleep may lead to an overall better retention of the exemplar value space. To investigate this, we predicted participants’ choices in the implicit memory generalization test using a feature selection approach based on the individual weights assigned to the different relevant and irrelevant feaures (see Fig. 2E).

First, we predicted performance in the implicit memory test using only the three features relevant for classification, disregarding the irrelevant feature. As expected, this analysis replicated the results from the overall model prediction that included the irrelevant feature, confirming a better transfer of the acquired value space for the relevant features in the night-sleep group compared to the day-wake group, while no circadian differences were observed (night-sleep [66.89 % ± 0.77 %] ≠ day-wake group [63.90 % ± 0.97 %]: *t*(124) = 2.41, *p* = .017, *d* = 0.43; control morning [63.93 % ± 1.67 %] ≠ control evening group [66.28 % ± 1.21 %]: *t*(55) = −1.14, *p* = .258, *d* = −0.30; Fig. 6A; Supplementary Tables S6A, S7A). We then tested if sleep selectively benefits a combined, multi-dimensional feature use compared to merely strengthening memory for single features. For this, we predicted participants’ implicit memory performance with each individual relevant feature separately and subtracted the resulting averaged single-relevant-feature prediction accuracy from the combined three-relevant-features prediction accuracy. However, this analysis only yielded a descriptive benefit of the night-sleep compared to the day-wake group in an incremental combined use of the relevant features compared to only using single features that failed to reach significance (night-sleep [6.93 % ± 0.47 %] ≠ day-wake group [5.90 % ± 0.52 %]: *t*(124) = 1.45, *p* = .150, *d* = 0.26; Fig. 6B; Supplementary Tables S6B, S7B). No circadian differences were found (control morning [6.07 % ± 0.99 %] ≠ control evening group [5.40 % ± 0.69 %]: *t*(55) = 0.56, *p* = .581, *d* = 0.15; Fig. 6B). Together, this suggests that the better transfer of the learned exemplar value representation in the night-sleep group cannot be attributed to the use of a combined, multi-dimensional feature representation and may rather result from a better general retention of the acquired value space.

**Fig. 6.**
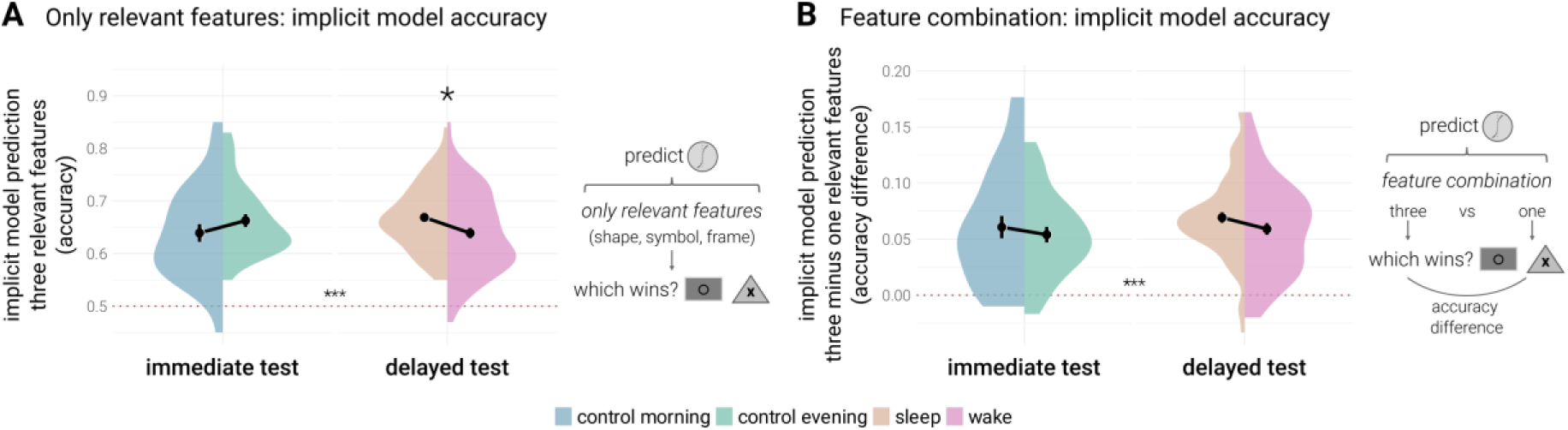
Model prediction in the implicit memory test based on selected features. (A) Only relevant features: Using only the three relevant features for prediction in the implicit memory test replicated the results from the overall model prediction with a significant benefit of the night-sleep compared to the day-wake group, while no circadian differences were observed. (B) Feature combination: Comparing the combined accuracy using all three relevant features with only one relevant feature did not yield significant differences between groups, suggesting that the better transfer of the learned exemplar value representation cannot be attributed to a stronger reliance on a combined, multi-dimensional feature representation. (A to B) Data are *M* ± *SEM*. Red dotted lines indicate chance performance (with asterisks above line indicating significant difference from chance performance), a value of 1 indicates 100 % accuracy. ***: *p* < .001; *: *p* < .05.

### Sleep does not promote explicit insight into the exemplar value rule

Participants in the night-sleep group compared to the day-wake group were more accurate in discriminating unseen exemplars based on their assigned value in the implicit memory test. We therefore tested whether participants in the sleep group also obtained better explicit insight into the task rule assigning values to exemplars. Participants were asked to indicate which factors determined exemplar value during the experimental task in written reports. These were rated based on whether the reported rules implied a hierarchy of exemplars or exemplar features, and based on whether the reported rules implied that a combination of exemplar features across exemplar dimensions defined exemplar value (nominal ratings yes/no). Although participants showed improved implicit memory performance after a night of sleep compared to a day of wakefulness, this did not transfer to enhanced explicit insight into the exemplar value rule, neither for rules involving a hierarchy (percentage of reported hierarchy rules: night-sleep group 54 %, day-wake group 54 %; Χ^2^(1) = 0.00, *p* = 1.000; Supplementary Tables S11D, S11E) nor for rules involving a combination of features (percentage of reported combination rules: night-sleep group 59 %, day-wake group 46 %; Χ^2^(1) = 1.56, *p* = .212; Supplementary Tables S11D, S11F). Although sleep allowed for better implicit generalization of the feature rule, this improvement did thus not transfer into the explicit, verbalizable domain.

### Modulation of memory-related brain activity by sleep

Analyzing differences in brain activity during memory tests between the night-sleep and day-wake groups did not yield any significant results at a conventional significance threshold (*p*_FWE_ < .05). Since we observed strong and reliable effects in behavioral outcomes for night-sleep compared to day-wake group comparisons, we report exploratory analyses at a liberal significance threshold in the Supplementary Information (*p*_uncorr_ < .005). Visualizations of all clusters and tables reporting all sleep > wake contrasts can be found in the Supplementary Information, including results on the explicit memory test where no behavioral benefit of sleep was observed (Supplementary Fig. S10; Supplementary Tables S8A-C, S9).

## Discussion

Interactions between memory systems are considerably more flexible than previously assumed. Here, we find that the relationship between explicit exemplar and implicit information-integration memory in a classification learning task is moderated by sleep. Sleep is known to support memory consolidation in multiple memory systems ^16,21,58^. Our data show that during sleep, memory is not only strengthened within these systems. Moreover, an initially competitive relationship between implicit and explicit memory is resolved after a night of sleep, but not after a day of wakefulness. This change in memory systems interaction is associated with a better transfer of the learned exemplar value representation to post-sleep memory tests assessing implicit memory for the exemplar hierarchy rule. Moreover, participants who slept showed superior performance both on the implicit memory generalization test, requiring use of implicit rule knowledge to assess novel exemplars not seen during classification task training, as well as on the known exemplar categorization task, where they could rely on both the explicit and implicit memory representations acquired during training. Sleep thus selectively helps to form implicit and explicit memory representations which may improve flexible strategy use depending on current task demands and aid generalization to novel situations.

Classification tasks, such as the one used in the present experiments, can be learned using explicit or implicit decision strategies and, accordingly, have been shown to activate the medial temporal lobe or the striatum, depending on boundary conditions such as training mode, training duration, or the activation of the stress system ^8,9,59–62^. Moreover, activity in multiple memory systems can shift over time, possibly reflecting strategy use ^63^. The striatum and medial temporal lobe can also be recruited in parallel in other memory tasks, such as motor sequence learning, associative memory, (reward) generalization, and navigation ^10,55,56,64–69^. Here, explicit and implicit learning systems can interact both competitively and cooperatively. Sleep has been proposed as an important factor modulating the nature of this interaction ^11,70^. Our task was designed to examine the development of explicit and implicit memory components and their interaction. We show that, after sleep, the initially competitive interaction between explicit and implicit memory is resolved. A mutual inhibition, reflected in a negative relationship between these two aspects of memory after training, was replaced by a non-competitive relationship after sleep. Importantly, sleep also benefitted performance in a memory test allowing for the use of both explicit and implicit memory systems. Sleep thus is a decisive factor in allowing for a flexible interplay between independent memory representations.

Our finding that sleep aids generalization of the acquired exemplar value rule to new exemplars extends research on qualitative changes in memory over sleep, including improved gist abstraction and extraction of hidden regularities ^40,43,46–48^. Note, however, that other studies failed to replicate this facilitating effect of sleep on the abstraction and generalization of task regularities ^34,71–73^. Our findings suggest that sleep may particularly favor implicit forms of generalization. Benefits of sleep on the extraction of hidden regularities in a wide range of learning tasks, including artificial grammar learning, information-integration learning, and category learning, were only evident in implicit performance improvement, but not explicit rule knowledge (note that this may differ for tasks employing temporal rules) ^74^. Consequently, participants’ performance on our implicit memory test benefitted from a night of sleep although they did not gain more explicit insight into the exemplar value rule.

While conclusions on sleep-dependent memory consolidation are usually drawn from direct measures of immediate and delayed memory performance, few attempts have been made to characterize the underlying cognitive representations using computational modeling approaches ^75^. We used a reinforcement learning approach to model participant-specific cognitive representations of the multi-dimensional feature space in an information-integration classification learning task, which could be solved by different mnemonic strategies. This allowed us to assess differences in strategy use between participants who slept after task training and participants who stayed awake. We find that after a night of sleep, participants made more efficient use of the exemplar value representation they had acquired during training. This suggests that sleep not only allows for better access to simple memory representations acquired before sleep, as typically indicated by better retention of learning material in declarative memory tasks ^18^, but may also enhance access to more complex representations of multidimensional task spaces ^76^. This greater cognitive availability of the task space may explain why participants who slept after training demonstrated superior generalization of the hidden exemplar value rule to novel, unseen exemplars, as reflected in their superior performance on the implicit memory generalization test. However, we did not find empirical support for the idea that sleep selectively benefits the use of a combined, multidimensional feature representation, as indicated by our feature selection analysis. Instead, sleep generally seems to help to access the acquired value space. Furthermore, whether the sleep-dependent benefit on generalization performance is driven by additional extraction or abstraction of the task rule from learned exemplars or reflects better retention of the rule acquired before sleep thus remains an open question.

Contrary to robust reports on sleep-dependent benefits in episodic memory performance ^58^, we did not find a beneficial effect of sleep on explicit exemplar recognition. The task goal in our study was to identify a hidden underlying exemplar hierarchy, regardless of whether participants employed an explicit exemplar-based or implicit information-integration strategy. Previous research has suggested that sleep-dependent memory consolidation mainly targets features with future relevance ^25,26^ (but see also ^27^). Consolidating individual exemplar representation may not have been of highest priority in the present experiments. Instead, sleep accomplished an implicit understanding of the exemplar hierarchy based on the explicitly learned exemplars and implicit information-integration rule representations before sleep. In line with task demands, this boosted implicit memory performance and led to resolved competition between explicit and implicit memory representations.

During sleep, the hippocampus coordinates reactivation of previous learning experiences together with other cortical and subcortical areas ^77–83^. Joint reactivation of memory representations in multiple memory systems may mechanistically underlie our finding that sleep resolved competition between implicit and explicit learning systems. Lansink et al. ^79^ investigated replay of neuronal activity during sleep in both the hippocampus and striatum. They show that firing patterns in which hippocampal cells code for the spatial component of the memory and striatal cells code for the associated reward component of the learning experience are preferentially reinstated in a coordinated replay across these brain structures. This cross-regional replay may have a functional relevance for consolidating a multi-facetted memory engram, with both spatial and reward components bound together during sleep-dependent consolidation. Our brain imaging analyses lacked the power to investigate neural evidence for such an integrated, multi-facetted memory representation. Behavioral evidence on the promotion of an integrated memory trace following reactivation is provided by a study using targeted-memory reactivation (TMR) during sleep indicating that reactivation promotes explicit knowledge of an implicit motor sequence rule ^84^. Furthermore, in a category learning paradigm, cueing of categories during sleep reduced details of the studied exemplars ^32^, indicating a reactivation-dependent rule abstraction. Our study extends these findings by demonstrating a sleep-dependent conversion of an initially competitive interaction between explicit and implicit memory systems following sleep, assessed via behavioral performance. During initial task training, separate memory representations are formed in explicit and implicit memory systems. Over sleep, joint reactivation may lead to an integration of information across these systems. As a consequence, competing representations of the same task can be bound into one coherent representation, so that different aspects of a memory can independently be accessed to contribute to optimal behavioral output. This is reflected in a non-competitive relationship between explicit and implicit memory performance, as well as better performance on known exemplar categorization which can be solved using both explicit and implicit memory representations after sleep.

Our findings demonstrate that sleep plays a critical role in resolving competition between explicit exemplar and implicit information-integration memory systems, fostering independent representations that enhance memory performance and generalization. By resolving competition between memory representations, sleep enables the formation of a more coherent and accessible memory trace, improving implicit rule abstraction and generalization in an information-integration category learning task and the application of learned information to novel situations. These results align with prior research on qualitative changes to memory over sleep-dependent consolidation and extend these findings by highlighting that sleep affects the flexible interplay between multiple memory systems in addition to strengthening individual memory traces. While our behavioral data suggest that sleep promotes cross-systems strategy deployment, future neuroimaging studies are needed to directly assess the neural mechanisms underlying this process. Overall, our study provides compelling evidence that sleep not only stabilizes memory but also transforms the underlying cognitive architecture, promoting the adaptive use of stored knowledge.

## Methods

### Participants

The main study followed a 12-h day-wake vs. 12-h night-sleep design (see Fig. 1A). 87 participants (52 female, 35 male) learned a feedback-driven classification task either in the morning 1 hour after their habitual wake-up time (*n* = 44, day-wake group), or in the evening after being awake for 13 hours (*n* = 43, night-sleep group). Performance was tested after a 12-h consolidation period, during which they had either a full night of sleep or a normal day of wakefulness. All participants were healthy adults between 18 and 30 years with no history of neurological or psychiatric disorders. They were color-sighted as assessed by the Ishihara test. They were regular sleepers with a habitual sleep duration of more than 6 hours and wake up times between 7 am and 9 am. They did not report any chronic or acute sleep related problems, did no shift work, and had not changed time zones in the 6 weeks leading up to the experiment. Participants were told to refrain from drinking alcohol, coffee, and tea on the day of the experiment, as well as the night before, and did not take any medication that affect the central nervous system. All participants were non-smokers. Participants provided written informed consent to undergo all study procedures before the study commenced. All procedures were approved by the ethics committee at the LMU Munich (Department of Psychology, Ludwig-Maximilians-Universität München, Germany, reference GA730/3-1). Participants were randomly assigned to either the sleep or wake condition, with sex balanced across conditions. Participants kept sleep logs during the two nights preceding the experimental session and on the day of the experiment itself to verify regular sleeping patterns. Additionally, activity during the consolidation period was monitored using actimetry to confirm that participants in the wake groups did not sleep during the day and to verify reports on sleep length in the sleep group. Sleep journals and actimetry data were found to be compatible and no participant was excluded for this reason. One participant from the night-sleep group who showed insufficient learning performance during the second half of training (< 70 % exemplars correctly learned) was excluded, resulting in a sample size of *n* = 42 for the night-sleep group. Participants in the night-sleep group slept on average 7 h 9 min ± 8 min during the night after learning the classification task. At the end of each visit to the laboratory, a subset (*n* = 40) of participants performed a 10-min psychomotor vigilance task (PVT) ^85^. Fatigue did not differ between conditions (mean number of lapses – learning: sleep 1.76 ± 0.49, wake 0.95 ± 0.30; *t*(38) = 1.38, *p* = .177; retrieval: sleep 0.90 ± 0.26, wake 0.84 ± 0.32; *t*(38) = 0.15, *p* = .878; median reaction times – learning: sleep 295.26 ± 5.41 *ms*, wake 298.66 ± 5.43 *ms*; *t*(38) = −0.44, *p* = .661; retrieval: sleep 289.62 ± 5.73 *ms*, wake 295.61 ± 5.53 *ms*; *t*(38) = −0.75, *p* = .459).

In a second experiment designed to control for circadian influences, 58 participants (42 female, 16 male) learned the task again either in the morning (*n* = 29) or in the evening (*n* = 29) but were retested immediately after training to control for circadian effects on performance (Fig. 1A). One participant from the morning group who showed insufficient learning performance during the second half of training (< 70 % exemplars correctly learned) was excluded, resulting in a sample size of *n* = 28 for the morning control group. A third experiment with 40 participants (31 female, 9 male) followed the same day-wake/night-sleep design (*n* = 19 and *n* = 21, respectively) as the main study, but performance was tested in the magnetic resonance imaging (MRI) scanner (fMRI study, Fig. 1A). Participants in the night-sleep group slept on average 6 h 46 min ± 16 min during the night after learning the classification task. Sleep duration did not differ significantly between participants that were tested in the MRI scanner and participants that only performed the experimental task without MRI scanning (*t*(60) = 1.52, *p* = .133).

### Task

During classification task training, two stimuli were presented on a black screen, and participants had to decide which of these exemplars has a higher rank, i.e., wins the comparison (Fig. 1C). The rank of the stimuli was defined by a hidden rule reflecting a linear combination of values assigned to stimulus properties, which is typical for information-integration tasks ^86,87^. Stimuli differed on four dimensions (shape, symbol, fill color, frame color).

Each dimension had five possible properties (e.g., triangle, ellipse, rectangle, square, circle for the dimension ‘shape’). For each participant, we assigned a new arbitrary hierarchy to the five properties of the dimensions shape, symbol, and fill color (exemplary hierarchy shown in Fig. 1B). According to the hierarchy, scores between 1 and 5 were assigned to the properties. Frame color served as a distractor. A score between 3 and 15 was calculated for each object by adding these values across dimensions (Fig. 1E). Because of this small range of scores for the 625 possible different stimuli, this task was difficult to learn. Together with the irrelevant distractor dimension, it was impossible for the subjects to deduce the full underlying value system explicitly. When asked whether they could name general rules that determined exemplar values at the end of the experiment, participants were unable to explicitly report the complete information-integration rule.

During training, participants’ task was to decide which of the two presented exemplars ranked higher according to this score, and to indicate the result via keypress with the left or right index finger, respectively (Fig. 1C). They performed 625 of these decision trials. In each trial, the two exemplars were shown for a minimum of 2 s. If participants did not react within 5 s, the stimuli vanished, and a black screen was shown until a decision was made. The short presentation time and the large number of trials made it virtually impossible to keep all eight different features of one display in working memory. We intended this procedure to prevent participants from actively trying to discover the hidden rule underlying the classification task by intensely studying, comparing, and excluding different alternatives based on the stimuli present on screen. After each reaction, a symbolized happy or sad face was presented as feedback for 2 s, depending on whether the answer was correct or not. If the exemplars had already disappeared when the decision was made, they reappeared together with the smiley. The sad face was additionally accompanied by an aversive sound and followed by an empty screen and a penalty delay of 3 s, emphasizing the error.

During the 625 training trials, 50 exemplars were shown in varying combinations in such a way that 25 exemplars were always the higher-ranking ones (winner exemplars; exemplar sum point value between 10 and 12) and 25 exemplars were always the lower-ranking ones (loser exemplars; exemplar sum point value between 6 and 8; Fig. 1E). Thus, participants could solve the task either by *explicitly* remembering for single exemplars if they were winners or losers (*exemplar-based* decision), or by *implicitly* learning and generalizing the underlying hidden regularities that determined the exemplar’s rank (decision based on *information-integration* learning). Instructions made participants aware of both possible solutions. They were also informed that they would be asked to recognize the stimuli later.

In the retrieval session, three different aspects of memory were tested (see Fig. 1D). Implicit memory for the hidden regularities was tested by pairing two new exemplars (exemplar sum point values between 3 and 15; Fig. 1E). On each of the 50 trials, participants had to decide which exemplar ranked higher. These decisions could only be based on implicit memory for the underlying hidden rule acquired during information-integration learning. Explicit exemplar memory was tested in a recognition task by presenting those 50 exemplars that had appeared during learning and 50 new ones. Participants indicated whether an exemplar was old or new (total of 100 trials). To successfully solve this task, participants had to use their explicit memory of the exemplars presented during training. Additionally, we used a categorization task that could be solved with the help of both explicit and implicit memory. On each of the 50 trials, participants were asked for individual, previously presented exemplars to judge whether these exemplars would rather win or lose in comparison to other exemplars. In this task, best performance can be achieved by relying on both explicit and implicit memory representations (explicit: correctly remember exemplar value; implicit: correctly infer exemplar value from acquired rule representation). Finally, in a forced-choice task, we created a competition between explicit exemplar memory and memory for the implicit information-integration rule. The 25 winner exemplars and 25 loser exemplars from training were paired with higher-ranking (sum point values of 11 to 15) and lower-ranking new items (sum point values of 3 to 7), respectively, such that winner exemplars would now lose, and loser items would now win the comparison (total of 50 trials). This task was designed to assess relative changes in the use of explicit and implicit memory representations. We found that participants tended to rely on exemplar-based memory in this forced-choice task, as indicated by relatively more exemplar-based (explicit) compared to rule-based (implicit) decisions, yet no differences emerged between conditions. Results on this task are reported in the Supplementary Information (Supplementary Tables S2E, S3E). During all retrieval tests, no feedback for correctness of decisions was given. The order of memory tests was the same for each participant, starting with the forced-choice task, then the explicit and known exemplar categorization memory tests, and finishing with the implicit memory test. Finally, participants’ explicit insight into the underlying task rule was assessed by asking participants to indicate what determined exemplar value in the experimental task using written reports.

The task design was chosen based on a large literature of studies on classification learning ^86^. It has been shown that various strategies are used to solve classification tasks ^60^. We specifically enforced multiple systems use by providing strong cues for different learning strategies during task training. For one, we prompted implicit information-integration learning by basing the hidden rule on a conjunction of features, which is in its complexity impossible to verbalize and has to be inferred from integrating information across many stimuli and feature dimensions. This kind of implicit learning has been shown to depend critically on the striatum ^6,7^. Feedback-driven training is thought to further reinforce these implicit aspects of memory and striatal involvement ^8,88,89^. The use of explicit exemplar-based strategies was ensured by instructing subjects about the possibility to classify each exemplar as belonging to the winning or to the losing category. During classification learning, every exemplar can be stored in memory, along with its winning or losing category label. This kind of explicit encoding strategy depends critically on the hippocampus ^8,90^.

In the fMRI experiment, the retrieval tests and retrieval test timings were equivalent to behavioral experiment. Every 60 s, all tests were interrupted by short blocks of an odd-even number judgement task which has been shown to suppress hippocampal activity ^91^. Block duration for the odd-even judgement task was jittered between 18 and 23 s, with an average duration of 20.5 s. Participants had 1.5 s per trial to select their response, and each trial was followed by a 0.5 s intertrial interval.

### Behavioral data analysis

Behavioral analyses were performed on the collapsed behavioral data from the main (no MRI scanning) and the fMRI study. Analyses conducted separately for the main and the fMRI study can be found in the Supplementary Information. Descriptive statistics are provided as mean values (± standard error). All statistical comparisons were two-tailed tests at a significance level of *α* = 0.05.

#### Memory test scores

Initial learning performance was measured as the percentage of correct decisions during the second half of classification task training. Implicit information-integration memory was measured as the percentage of correct decisions in the implicit memory test. Explicit exemplar recognition was analyzed in terms of memory sensitivity *d*’ which is calculated as *Z*(ℎ*its*) − *Z*(*false alarms*) in the recognition task. Known exemplar categorization was assessed by the percentage of correct decisions whether items would rather win or lose in the categorization task. Chance performance was at 50 % for the implicit memory test and the categorization task. Memory systems competition was measured as percentage of rule-based decisions. Chance performance cannot be meaningfully determined in this forced-choice test.

#### Relationship between explicit and implicit memory test scores

We analyzed the relationship between explicit and implicit aspects of memory using multiple regression analysis. We corrected for initial learning performance when testing relationships between memory scores at test by regressing out performance in the second half of initial learning before entering memory scores into the final regression analysis to account for differences in initial learning level. The explicit memory score and experimental group (night-sleep, day-wake, morning control, and evening control) served as predictors while the implicit memory score served as dependent variable. We examined regression slopes for the individual groups and compared slopes between experimental groups. As such, we investigated differences in the relationship between explicit and implicit memory for participants who slept after learning and participants who stayed awake (main and fMRI study). We then compared this to differences in the relationship between explicit and implicit memory for participants that were directly tested on their explicit and implicit memory after learning either in the morning or in the evening (circadian control groups).

#### Group differences in memory test scores

We compared differences in memory performance (initial learning performance, implicit information-integration memory, explicit exemplar recognition, and known exemplar categorization) between experimental groups using Student’s *t*-tests. Performance in the second half of initial learning was regressed out before entering memory test scores into the *t*-tests to account for differences in initial learning level. Note that in the figure (Fig. 4) accompanying this analysis, uncorrected memory scores are displayed to aid comprehension although statistical tests were performed on residual scores after learning performance in the second half of learning has been regressed out. Cohen’s *d* is reported as effect size.

#### Explicit insight into exemplar value rule

Participants’ written reports on what determined exemplar value during the experimental task were rated for indications of hierarchy and combination rules (nominal ratings yes/no). A self-reported rule was rated as a hierarchy rule if a report either explicitly or implicitly referred to a hierarchy between exemplars or exemplar features. A self-reported rule was rated as a combination rule if a report mentioned the combination of exemplar dimensions defining exemplar value. Rules were scored by two independent raters. Rater agreement for the hierarchy rule was at 88 % and for the combination rule at 96 % (see Supplementary Table S11A). If the two raters did not agree on the same result, no indication of a reported hierarchy or combination rule was scored in the final rating. Validity of the applied rating approach is suggested by improved performance in the implicit memory test in participants reporting a hierarchy or combination rule (see Supplementary Tables S11B, S11C). Differences between the number of reported rules between groups were assessed using Χ^2^-tests.

### Reinforcement learning (RL) model

Participants’ behavior during task training and memory tests was modeled using a reinforcement learning (RL) algorithm (see Fig. 2B) to assess participant-specific exemplar value representations. The goal of this analysis was to test to which extent they applied resulting exemplar (explicit) and value (implicit) representations during later memory tests by predicting participants’ behavior based on the learned model representations.

#### Environment formulation

We formulate the classification task as a contextual multi-armed bandit task with action space *A*. Specifically, we define each action (exemplar) *a* ∈ *A* as a vector consisting of three relevant feature dimensions (*shape*, *symbol*, *fill color*) and one irrelevant feature dimension (*frame color*). Each dimension may take one of five alternative values and their identity is randomized across participants, i.e., *a* ∈ ℤ^5^ and 1 ≤ *a* ≤ 5 for all *a* ∈ *A*. Thus, the entire action space has a cardinality of 5^4^ = 625.

#### Learning task and model fitting

We employ a reinforcement learning algorithm that is based on the assumptions made in the Rescorla-Wagner-model ^92,93^. (1) We assume that during the learning task, the action-values (*Q*-values) for the individual exemplars start with *Q*_0_(*a*) = 0 for all *a* ∈ *A*, and are then updated after the action execution for the *t*th trial according to:

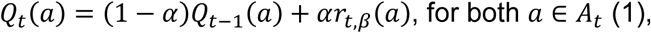

where *α* is the learning rate and *A_t_* is the action subspace of the *t*th trial, consisting of the presented left (*a*_L_) and right (*a*_R_) exemplars, i.e., *_At_* = {*a*_L_, *a*_R_}. (2) Note that we define positive and negative reward values *rt*_,*β*_ for the winner and loser exemplars, respectively, according to

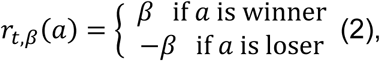

where *β* is the reward strength. (3) We then apply a Boltzmann policy with respect to the action-value function for the *t*th trial to select actions for each learning trial, i.e., given the action subspace *A_t_* = {*a*_L_, *a*_R_}, we have

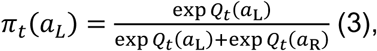

where *π_t_*(*a*_L_) describes the probability of selecting the left choice-option *a*_L_ over the right *a*_R_. (4) We use the data from the learning task to fit the algorithm parameters *α* and *β* for each individual participant. We then use these parameters to estimate the action-values in each trial of the subsequent test tasks. Specifically, we apply Markov chain Monte Carlo (MCMC) ^94^, a class of sampling-based parameter estimation methods. Let *D* be the participant behavior observed during the learning task, we aim to find the posterior distribution of model parameters *α* and *β* given *D*, i.e.,

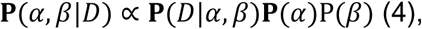

where we define the priors for *α* and *β* as *α* ∼ Beta(2,5) and *β* ∼ Ga(7.5,1), respectively. To sample from the posterior **P**(*α*, *β*|*D*), we sample 4 Markov chains with 5,000 samples in parallel (after 2,000 discarded burn-in samples) drawn from each chain. Afterwards, we select the group of parameters with highest posterior probability (i.e., the posterior mean) for further analysis. As a result, we obtain an exemplar value representation of the exemplars studied during task training, represented by participants’ individual *Q*-values.

#### Generalization of action-values

To predict choices in the test tasks, we generalize the learned action-values of familiar, previously seen exemplars to new, unseen exemplars. (5) To this end, we assume that

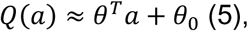

for all *a* ∈ *A*, where *θ* ∈ ℝ^4^ describes the coefficient vector of the four feature dimensions of each exemplar and *θ*_0_ ∈ ℝ denotes the intercept. (6) For each participant, we estimate the parameters *θ* and *θ*_0_ by solving the following least-squares problem:

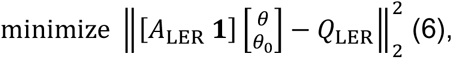

where *A*_LER_ ∈ ℤ^50×4^ describes the action subspace of the familiar exemplars that were previously seen during learning (LER) and *Q*_LER_ describes the final action-value function over *A*_LER_ after learning.

#### Test tasks

For each of the three test tasks, we then model the decision policies according to the instructions for the individual tasks.

##### Explicit memory task

(7) For the *explicit memory* task, we employ a feature-based decision policy to model the probability of a participant to indicate that they have seen the exemplar before (*true*):

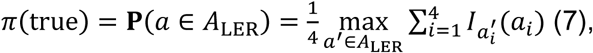

where 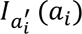 is an indicator function that equals to 1 if 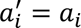 and 0 otherwise. This policy works such that the probability to label an exemplar as old (instead of new) in the recognition memory test is higher the more similar a presented exemplar is to an exemplar that was actually presented during task training.

##### Implicit memory task

(8) For the *implicit memory* task all *a* ∉ *A*LER. Thus, we employ a Boltzmann policy on the generalized action-values:

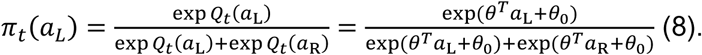

Therefore, this policy makes use of the exemplar value representation acquired during task training (*Q*-values) and generalizes them to decisions between exemplars that were never presented before.

##### Categorization task

(9) For the *categorization* task where all *a* ∈ *A*_LER_, we employ the following policy to estimate the probability that the participant would identify a given exemplar *a* as a winner (*win*):

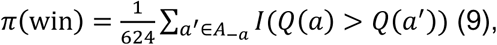

where *A*−*a* is the action space with the current test exemplar *a* removed, and the indicator function *I*(*Q*(*a*) > *Q*(*a*^′^)) is equal to 1 if *Q*(*a*) > *Q*(*a*^′^) and 0 otherwise, with the generalized action-values *Q*(*a*) = *θ^T^_a_* + *θ*_0_ for *a* and all *a*^′^ ∈ *A*_−*a*_. This policy is more likely to label a presented exemplar as winner (compared to loser) the higher the exemplar ranks among all possible exemplars and therefore makes use of both exemplar value representations acquired during task training and generalized exemplar value representations of exemplars not presented during training.

#### Feature selection analysis

Following prediction using all features in the implicit memory task, a features selection analysis was performed in which only selected features were used for prediction of implicit memory performance. Resulting values indicate the degree to which the exemplar value representation acquired during training for the selected feature or combination of features was transferred to later memory tests.

### Functional MRI acquisition and analysis

Whole-brain functional T2*- weighted MRI data were acquired at the Max Planck Institute of Psychiatry, Munich, with a 3-T MR750 scanner (GE) using a 12-channel head coil and a single-shot echo planar imaging (EPI) sequence (repetition time: 2,500 ms; echo time: 30 ms; flip angle: 90°; voxel size: 1.88 x 1.88 x 3.5 mm^3^, acquisition matrix size: 96 x 96 x 42, reconstructed to 128 x 128 x 42 voxel; interleaved slice acquisition). A whole brain structural scan was acquired using a T1-weighted 3D fast spoiled gradient echo sequence (repetition time: 7.1 ms; echo time: 2.2 ms; flip angle: 12°; phase acceleration: 2; voxel size: 0.8 x 0.8 x 1.3 mm^3^; matrix size: 320 x 320 x 128 voxel). DICOM to NIFTI conversion was performed using `dcm2niix` ^95^. The first four scans were discarded to allow for magnetic saturation effects. Results included in this manuscript come from preprocessing performed using *fMRIPrep* 23.2.0 (RRID:SCR_016216) ^96,97^, which is based on *Nipype* 1.8.6 (RRID:SCR_002502) ^98,99^.

#### Anatomical data preprocessing

The T1w image was corrected for intensity non-uniformity (INU) with `N4BiasFieldCorrection` ^100^, distributed with ANTs 2.5.0 (RRID:SCR_004757) ^101^, and used as T1w-reference throughout the workflow. The T1w-reference was then skull-stripped with a *Nipype* implementation of the `antsBrainExtraction.sh` workflow (from ANTs), using OASIS30ANTs as target template. Brain tissue segmentation of cerebrospinal fluid (CSF), white-matter (WM) and gray-matter (GM) was performed on the brain-extracted T1w using `fast` (FSL, RRID:SCR_002823) ^102^. Brain surfaces were reconstructed using `recon-all` (FreeSurfer 7.3.2, RRID:SCR_001847) ^103^, and the brain mask estimated previously was refined with a custom variation of the method to reconcile ANTs-derived and FreeSurfer-derived segmentations of the cortical gray-matter of Mindboggle (RRID:SCR_002438) ^104^. Volume-based spatial normalization to one standard space (MNI152NLin2009cAsym) was performed through nonlinear registration with `antsRegistration` (ANTs 2.5.0), using brain-extracted versions of both T1w reference and the T1w template. The following template was selected for spatial normalization and accessed with *TemplateFlow* (23.1.0) ^105^: *ICBM 152 Nonlinear Asymmetrical template version 2009c* (RRID:SCR_008796; TemplateFlow ID: MNI152NLin2009cAsym) ^106^.

#### Functional data preprocessing

For each of the 4 BOLD runs found per subject (across all tasks and sessions), the following preprocessing was performed. First, a reference volume was generated, using a custom methodology of *fMRIPrep*, for use in head motion correction. Head-motion parameters with respect to the BOLD reference (transformation matrices, and six corresponding rotation and translation parameters) are estimated before any spatiotemporal filtering using `mcflirt` (FSL) ^107^. The BOLD reference was then co-registered to the T1w reference using `bbregister` (FreeSurfer) which implements boundary-based registration ^108^. Co-registration was configured with six degrees of freedom. Several confounding time-series were calculated based on the *preprocessed* BOLD: framewise displacement (FD), DVARS and three region-wise global signals. FD was computed using two formulations following Power (absolute sum of relative motions) ^109^ and Jenkinson (relative root mean square displacement between affines) ^107^. FD and DVARS are calculated for each functional run, both using their implementations in *Nipype* (following the definitions by ^109^). The three global signals are extracted within the CSF, the WM, and the whole-brain masks. Six head-motion estimates (3 translations, 3 rotations) calculated in the correction step were placed within the corresponding confounds file. Frames that exceeded a threshold of 0.5 mm FD or 1.5 standardized DVARS were annotated as motion outliers. All resamplings were performed with a *single interpolation step* by composing all the pertinent transformations (i.e., head-motion transform matrices and co-registrations to anatomical and output spaces). Gridded (volumetric) resamplings were performed using `nitransforms`, configured with cubic B-spline interpolation.

Many internal operations of *fMRIPrep* use *Nilearn* 0.10.2 (RRID:SCR_001362) ^110^, mostly within the functional processing workflow. For more details of the pipeline, see the section corresponding to workflows in fMRIPrep’s documentation.

#### Copyright waiver

The above boilerplate text was automatically generated by *fMRIPrep* with the express intention that users should copy and paste this text into their manuscripts unchanged. It is released under the CC0 license. Sections referring to estimates not considered in our preprocessing workflow were omitted.

#### Preprocessing quality control

Quality control of anatomical and functional data preprocessing was ensured by visual inspection of fMRIPrep’s output summary. For each subject, brain mask and brain tissue segmentation of the T1w, spatial normalization of the anatomical T1w reference, and surface reconstruction were examined. Furthermore, FD, DVARS and the three region-wise global signals were checked for large deviations suggesting excessive motion. Indices of all subjects were found to suffice for subsequent data analysis.

#### General linear model analysis

Mixed-effects general linear model (GLM) analyses were performed using SPM12 (SPM, RRID:SCR_007037). Before data were entered into the GLM analyses, they were smoothed with an 8 mm FWHM Gaussian kernel.

##### Memory test functional brain activity

We assessed brain regions contributions to correct memory recall and classification performance in GLM analyses modeling brain activity during trials in which participants made correct versus incorrect decisions that differed between night-sleep and day-wake groups. At the first level, for each memory test, onsets and durations until motor responses following stimulus presentation (separately for correct decisions, incorrect decisions and responses to the odd-even number judgement task) were convolved with a standard hemodynamic response function. These were modeled together with 6 head-motion regressors obtained from fMRIPrep (three rotations and three translations) as fixed effects in a GLM for each individual subject. High-pass filtering was implemented in the matrix design by using a cutoff period of 128 sec to remove low-frequency drifts from the time series. Serial correlations in the signal were estimated by using a first-order autoregressive plus white noise model and a restricted maximum likelihood (ReML) algorithm. The effects of interest consisted in the hemodynamic response to correct and incorrect decisions in the different memory tests. This was tested by obtaining statistical parametric maps from positive contrasts for correct and incorrect decisions separately. These summary statistic images were further smoothed (6 mm FWHM Gaussian Kernel) and entered in a second-level analysis where data of all participants were entered in a full factorial random effects model with the factors performance (correct vs. incorrect decisions) and group (night-sleep vs. day-wake). Participants were treated as a random factor. Main effects and interactions were tested using *t*-contrasts. ReML estimates of variance components were used to allow for unequal variance and possible deviations from sphericity.

##### Q-value modulation

In GLM analyses on modulation by *Q*-value, correlations between *Q*-values obtained from the RL model and brain activity in the implicit memory test were performed. At the first level, a *Q*-value regressor was obtained by calculating the absolute difference between participants’ *Q*-values of the two exemplars in the present implicit memory test trial and convolving this with a standard hemodynamic response function. Therefore, this analysis tests for brain regions that display variance in their activity according to the difference between participants’ value representations of the presented unknown exemplars. Similar to the GLM analysis on modulation of brain activity according to memory test performance, the *Q*-value regressor was modeled together with responses to the odd-even number judgement task and 6 head-motion regressors obtained from fMRIPrep (three rotations and three translations) as fixed effects in a GLM for each individual subject. High-pass filtering was implemented with a cutoff period of 128 s, and serial correlations in the signal were estimated by using a first-order autoregressive plus white noise model and a restricted maximum likelihood (ReML) algorithm. The effect of interest consisted in the hemodynamic response to variations in the *Q*-value difference in the implicit memory test. This was tested by obtaining a statistical parametric map from this contrast. This summary statistic image was further smoothed (6 mm FWHM Gaussian Kernel) and entered in a second-level analysis testing for differences in *Q*-value modulation between groups (night-sleep vs. day-wake, two-sample *t*-test) treating participants as a random factor (see above).

##### Significance threshold

No significant differences between night-sleep and day-wake groups in functional brain activity were observed on conventional statistical thresholds (*p*_FWE_ < .05). Therefore, exploratory analyses on *p*uncorr < .005 were performed as we observed strong and reliable effects in behavioral outcomes but not in the corresponding GLM analyses on brain activity and are reported in the Supplementary Information (Supplementary Tables S8A-C, S9; Supplementary Fig. S10). All coordinates are given as standard Montreal Neurological Institute (MNI) coordinates and correspond to the maxima of the reported cluster of activation. Coordinates were labeled using `bspmview`. Functional images are displayed on a standard MNI T1 image.

## Acknowledgements

A special thanks to Eliana Feurer and Lena Tesch for handling and scoring the written rule reports. Also, thanks to Anika Löwe for helpful comments and discussion of the manuscript.

## Data and code availability

Behavioral data and supporting analysis code will be made available on OSF (Open Science Framework) upon publication.

## Author contributions

Florian Pargent, Steffen Gais, and Monika Schönauer conceived and designed the experiments. Florian Pargent, Jana Werle, Michael Czisch, Steffen Gais, and Monika Schönauer performed the experiments. Katja Kleespies, Philipp C. Paulus, Hao Zhu, Marie Jakob, and Monika Schönauer analyzed the data. Katja Kleespies, Philipp C. Paulus, Hao Zhu, Marie Jakob, Michael Czisch, Joschka Boedecker, and Monika Schönauer contributed materials/analysis tools. Katja Kleespies, Philipp C. Paulus, Hao Zhu, Steffen Gais, and Monika Schönauer wrote the paper.

## Competing interests statement

All authors declare no competing interests.

## Notes

### Competing Interest Statement

The authors have declared no competing interest.

### Summary of Updates

Updated behavioral analyses and interpretation; removed MRI results from main text.

