## Supplementary Information for "Sleep resolves competition between explicit exemplar memory and implicit memory generalization"

#### Supplementary Information 1: Relationships between explicit and implicit memory performance

***Table S1A.*** Relationship between explicit and implicit memory performance in behavioral (main and circadian control) study groups.

| group (comparison) | test type | outcome | $b$ | $SE$ | $t$ | $df$ | $p$ |
|---|---|---|---|---|---|---|---|
| control morning | immediate test | explicit*implicit | $-0.09$ | $0.03$ | $-3.23$ | $135$ | $.002$** |
| control evening | immediate test | explicit*implicit | $-0.09$ | $0.03$ | $-2.84$ | $135$ | $.005$** |
| wake | delayed test | explicit*implicit | $-0.09$ | $0.03$ | $-3.26$ | $135$ | $.001$** |
| sleep | delayed test | explicit*implicit | $0.01$ | $0.02$ | $0.54$ | $135$ | $.589$ |
| control morning ≠ control evening | immediate test | explicit*implicit | $0.00$ | $0.04$ | $-0.06$ | $135$ | $.952$ |
| sleep ≠ wake | delayed test | explicit*implicit | $0.11$ | $0.04$ | $2.90$ | $135$ | $.004$** |
| (control evening – control morning) ≠ (sleep – wake) | immediate vs delayed test | explicit*implicit | $-0.11$ | $0.06$ | $-1.97$ | $135$ | $.051$ |

*Note*. **: $p < .01$.

***Table S1B.*** Relationship between explicit and implicit memory performance in fMRI study groups.

| group (comparison) | test type | outcome | $b$ | $SE$ | $t$ | $df$ | $p$ |
|---|---|---|---|---|---|---|---|
| wake | delayed test | explicit*implicit | $-0.09$ | $0.03$ | $-2.72$ | $36$ | $.010$* |
| sleep | delayed test | explicit*implicit | $0.05$ | $0.04$ | $1.13$ | $36$ | $.268$ |
| sleep ≠ wake | delayed test | explicit*implicit | $0.14$ | $0.05$ | $2.61$ | $36$ | $.013$* |

*Note*. *: $p < .05$.

***Table S1C.*** Relationship between explicit and implicit memory performance in combined behavioral (main and circadian control) and fMRI study groups.

| comparison | test type | outcome | $b$ | $SE$ | $t$ | $df$ | $p$ |
|---|---|---|---|---|---|---|---|
| control morning | immediate test | explicit*implicit | $-0.09$ | $0.03$ | $-3.25$ | $175$ | $.001$** |
| control evening | immediate test | explicit*implicit | $-0.09$ | $0.03$ | $-2.91$ | $175$ | $.004$** |
| wake | delayed test | explicit*implicit | $-0.08$ | $0.02$ | $-3.55$ | $175$ | $< .001$*** |
| sleep | delayed test | explicit*implicit | $0.01$ | $0.02$ | $0.74$ | $175$ | $.460$ |
| control morning ≠ control evening | immediate test | explicit*implicit | $0.00$ | $0.04$ | $-0.10$ | $175$ | $.923$ |
| sleep ≠ wake | delayed test | explicit*implicit | $0.09$ | $0.03$ | $3.09$ | $175$ | $.002$** |
| (control evening – control morning) ≠ (sleep – wake) | immediate vs delayed test | explicit*implicit | $0.09$ | $0.05$ | $1.88$ | $175$ | $.062$ |

*Note*. ***: $p < .001$; **: $p < .01$.

### Supplementary Information 2: Group summaries of task behavioral performance

***Table S2A.*** Learning (2nd half) behavioral performance group summary.

| group | study type | $M$ | $SE$ |
|---|---|---|---|
| control morning | main | 91.98 | 1.38 |
| control evening | main | 92.53 | 0.93 |
| sleep | main | 91.84 | 0.94 |
| wake | main | 92.56 | 0.95 |
| overall | main | 92.23 | 0.52 |
| sleep | fMRI | 90.95 | 0.74 |
| wake | fMRI | 91.61 | 1.53 |
| overall | fMRI | 91.26 | 0.81 |
| control morning | combined | 91.98 | 1.38 |
| control evening | combined | 92.53 | 0.93 |
| sleep | combined | 91.54 | 0.67 |
| wake | combined | 92.27 | 0.80 |
| overall | combined | 92.02 | 0.44 |

***Table S2B.*** Explicit behavioral performance group summary.

| group | study type | $M_{res}$ | $SE_{res}$ | $M$ | $SE$ |
|---|---|---|---|---|---|
| control morning | main | 0.22 | 0.09 | 1.12 | 0.10 |
| control evening | main | 0.00 | 0.07 | 0.91 | 0.08 |
| sleep | main | −0.04 | 0.07 | 0.86 | 0.08 |
| wake | main | −0.11 | 0.05 | 0.81 | 0.06 |
| overall | main | 0.00 | 0.04 | 0.90 | 0.04 |
| sleep | fMRI | 0.05 | 0.07 | 0.76 | 0.07 |
| wake | fMRI | −0.05 | 0.09 | 0.65 | 0.10 |
| overall | fMRI | 0.00 | 0.06 | 0.71 | 0.06 |
| control morning | combined | 0.26 | 0.09 | 1.12 | 0.10 |
| control evening | combined | 0.04 | 0.07 | 0.91 | 0.08 |
| sleep | combined | −0.03 | 0.05 | 0.82 | 0.06 |
| wake | combined | −0.11 | 0.05 | 0.76 | 0.05 |
| overall | combined | 0.00 | 0.03 | 0.86 | 0.03 |

***Table S2C.*** Implicit behavioral performance group summary.

| group | study type | $M_{res}$ | $SE_{res}$ | $M$ | $SE$ |
|---|---|---|---|---|---|
| control morning | main | −0.01 | 0.02 | 63.79 | 1.68 |
| control evening | main | 0.01 | 0.01 | 66.34 | 1.22 |
| sleep | main | 0.01 | 0.01 | 66.40 | 0.99 |
| wake | main | −0.01 | 0.01 | 64.07 | 1.17 |
| overall | main | 0.00 | 0.01 | 65.16 | 0.62 |
| sleep | fMRI | 0.02 | 0.01 | 67.67 | 1.17 |
| wake | fMRI | −0.02 | 0.02 | 63.53 | 1.73 |
| overall | fMRI | 0.00 | 0.01 | 65.70 | 1.06 |
| control morning | combined | −0.01 | 0.02 | 63.79 | 1.68 |
| control evening | combined | 0.01 | 0.01 | 66.34 | 1.22 |
| sleep | combined | 0.02 | 0.01 | 66.83 | 0.76 |
| wake | combined | −0.01 | 0.01 | 63.90 | 0.97 |
| overall | combined | 0.00 | 0.01 | 65.28 | 0.54 |

***Table S2D.*** Categorization behavioral performance group summary.

| **group** | **study type** | $M_{res}$ | $SE_{res}$ | $M$ | $SE$ |
|---|---|---|---|---|---|
| control morning | main | 0.01 | 0.01 | 81.29 | 2.02 |
| control evening | main | 0.02 | 0.01 | 83.52 | 1.63 |
| sleep | main | 0.01 | 0.01 | 81.76 | 1.14 |
| wake | main | −0.03 | 0.01 | 78.45 | 1.28 |
| overall | main | 0.00 | 0.01 | 81.01 | 0.74 |
| sleep | fMRI | 0.01 | 0.01 | 78.29 | 1.58 |
| wake | fMRI | −0.01 | 0.02 | 77.16 | 2.02 |
| overall | fMRI | 0.00 | 0.01 | 77.75 | 1.26 |
| control morning | combined | 0.01 | 0.01 | 81.29 | 2.02 |
| control evening | combined | 0.03 | 0.01 | 83.52 | 1.63 |
| sleep | combined | 0.01 | 0.01 | 80.60 | 0.94 |
| wake | combined | −0.02 | 0.01 | 78.06 | 1.07 |
| overall | combined | 0.00 | 0.00 | 80.30 | 0.65 |

***Table S2E.*** Competition behavioral performance group summary.

| **group** | **study type** | $M_{res}$ | $SE_{res}$ | $M$ | $SE$ |
|---|---|---|---|---|---|
| control morning | main | $-0.03$ | 0.02 | 35.43 | 2.12 |
| control evening | main | 0.00 | 0.02 | 37.93 | 2.64 |
| sleep | main | $-0.01$ | 0.02 | 37.52 | 1.65 |
| wake | main | 0.02 | 0.02 | 39.86 | 1.83 |
| overall | main | 0.00 | 0.01 | 37.92 | 1.00 |
| sleep | fMRI | 0.00 | 0.02 | 44.86 | 2.30 |
| wake | fMRI | 0.00 | 0.02 | 44.00 | 2.36 |
| overall | fMRI | 0.00 | 0.02 | 44.45 | 1.63 |
| control morning | combined | $-0.04$ | 0.02 | 35.43 | 2.12 |
| control evening | combined | $-0.01$ | 0.02 | 37.93 | 2.64 |
| sleep | combined | 0.00 | 0.01 | 39.97 | 1.40 |
| wake | combined | 0.02 | 0.01 | 41.11 | 1.47 |
| overall | combined | 0.00 | 0.01 | 39.34 | 0.88 |

**Supplementary Information 3: Group comparisons of task behavioral performance**

***Table S3A.*** Learning (2nd half) behavioral performance group comparisons.

| group (comparison) | study type | test type | outcome | $t$ | $df$ | $p$ | $d$ |
|---|---|---|---|---|---|---|---|
| control morning ≠ control evening | main | immediate test | learning accuracy (2nd half) | $-0.33$ | $55$ | $.740$ | $-0.09$ |
| sleep ≠ wake | main | delayed test | learning accuracy (2nd half) | $-0.54$ | $84$ | $.593$ | $-0.12$ |
| sleep ≠ control evening | main | evening training | learning accuracy (2nd half) | $-0.50$ | $69$ | $.616$ | $-0.12$ |
| wake ≠ control morning | main | morning training | learning accuracy (2nd half) | $0.36$ | $70$ | $.722$ | $0.09$ |
| overall ≠ chance performance | main | overall | learning accuracy (2nd half) | $81.92$ | $142$ | $< .001$*** | $6.85$ |
| sleep ≠ wake | fMRI | delayed test | learning accuracy (2nd half) | $-0.40$ | $38$ | $.690$ | $-0.13$ |
| overall ≠ chance performance | fMRI | overall | learning accuracy (2nd half) | $50.67$ | $39$ | $< .001$*** | $8.01$ |
| control morning ≠ control evening | combined | immediate test | learning accuracy (2nd half) | $-0.33$ | $55$ | $.740$ | $-0.09$ |
| sleep ≠ wake | combined | delayed test | learning accuracy (2nd half) | $-0.70$ | $124$ | $.487$ | $-0.12$ |
| sleep ≠ control evening | combined | evening training | learning accuracy (2nd half) | $-0.84$ | $90$ | $.403$ | $-0.19$ |
| wake ≠ control morning | combined | morning training | learning accuracy (2nd half) | $0.19$ | $89$ | $.847$ | $0.04$ |
| overall ≠ chance performance | combined | overall | learning accuracy (2nd half) | $95.41$ | $182$ | $< .001$*** | $7.05$ |

*Note*. ***: $p < .001$.

***Table S3B.*** Explicit behavioral performance group comparisons.

| comparison | study type | test type | outcome | $t$ | $df$ | $p$ | $d$ |
|---|---|---|---|---|---|---|---|
| control morning ≠ control evening | main | immediate test | explicit d’ | 1.93 | 55 | .058 | 0.51 |
| sleep ≠ wake | main | delayed test | explicit d’ | 0.82 | 84 | .417 | 0.18 |
| overall ≠ chance performance | main | overall | explicit d’ | 22.78 | 142 | < .001*** | 1.90 |
| sleep ≠ wake | fMRI | delayed test | explicit d’ | 0.88 | 38 | .384 | 0.28 |
| overall ≠ chance performance | fMRI | overall | explicit d’ | 12.21 | 39 | < .001*** | 1.93 |
| control morning ≠ control evening | combined | immediate test | explicit d’ | 1.90 | 55 | .063 | 0.50 |
| sleep ≠ wake | combined | delayed test | explicit d’ | 1.14 | 124 | .256 | 0.20 |
| overall ≠ chance performance | combined | overall | explicit d’ | 25.35 | 182 | < .001*** | 1.87 |

*Note*. ***: $p < .001$.

***Table S3C.*** Implicit behavioral performance group comparisons.

| comparison | study type | test type | outcome | $t$ | $df$ | $p$ | $d$ |
|---|---|---|---|---|---|---|---|
| control morning ≠ control evening | main | immediate test | implicit accuracy | $-1.20$ | $55$ | $.237$ | $-0.32$ |
| sleep ≠ wake | main | delayed test | implicit accuracy | $1.63$ | $84$ | $.106$ | $0.35$ |
| overall ≠ chance performance | main | overall | implicit accuracy | $24.39$ | $142$ | $< .001$*** | $2.04$ |
| sleep ≠ wake | fMRI | delayed test | implicit accuracy | $2.05$ | $38$ | $.048$* | $0.65$ |
| overall ≠ chance performance | fMRI | overall | implicit accuracy | $14.75$ | $39$ | $< .001$*** | $2.33$ |
| control morning ≠ control evening | combined | immediate test | implicit accuracy | $-1.20$ | $55$ | $.234$ | $-0.32$ |
| sleep ≠ wake | combined | delayed test | implicit accuracy | $2.51$ | $124$ | $.014$* | $0.45$ |
| overall ≠ chance performance | combined | overall | implicit accuracy | $28.43$ | $182$ | $< .001$*** | $2.10$ |

*Note*. ***: $p < .001$; *: $p < .05$.

***Table S3D.*** Categorization behavioral performance group comparisons.

| comparison | study type | test type | outcome | $t$ | $df$ | $p$ | $d$ |
|---|---|---|---|---|---|---|---|
| control morning ≠ control evening | main | immediate test | categorization accuracy | $-0.97$ | $55$ | $.335$ | $-0.26$ |
| sleep ≠ wake | main | delayed test | categorization accuracy | $2.75$ | $84$ | $.007$** | $0.59$ |
| overall ≠ chance performance | main | overall | categorization accuracy | $41.96$ | $142$ | $< .001$*** | $3.51$ |
| sleep ≠ wake | fMRI | delayed test | categorization accuracy | $0.97$ | $38$ | $.337$ | $0.31$ |
| overall ≠ chance performance | fMRI | overall | categorization accuracy | $22.11$ | $39$ | $< .001$*** | $3.50$ |
| control morning ≠ control evening | combined | immediate test | categorization accuracy | $-0.97$ | $55$ | $.336$ | $-0.26$ |
| sleep ≠ wake | combined | delayed test | categorization accuracy | $2.79$ | $124$ | $.006$** | $0.50$ |
| overall ≠ chance performance | combined | overall | categorization accuracy | $46.94$ | $182$ | $< .001$*** | $3.47$ |

*Note*. ***: $p < .001$; **: $p < .01$.

***Table S3E.*** Competition behavioral performance group comparisons.

| comparison | study type | test type | outcome | $t$ | $df$ | $p$ | $d$ |
|---|---|---|---|---|---|---|---|
| control morning ≠ control evening | main | immediate test | competition (rule-based) accuracy | $-0.90$ | $55$ | $.371$ | $-0.24$ |
| sleep ≠ wake | main | delayed test | competition (rule-based) accuracy | $-1.20$ | $84$ | $.234$ | $-0.26$ |
| overall ≠ chance performance | main | overall | competition (rule-based) accuracy | $-12.05$ | $142$ | $< .001^{***}$ | $-1.01$ |
| sleep ≠ wake | fMRI | delayed test | competition (rule-based) accuracy | $0.20$ | $38$ | $.846$ | $0.06$ |
| overall ≠ chance performance | fMRI | overall | competition (rule-based) accuracy | $-3.41$ | $39$ | $.002^{**}$ | $-0.54$ |
| control morning ≠ control evening | combined | immediate test | competition (rule-based) accuracy | $-0.90$ | $55$ | $.373$ | $-0.24$ |
| sleep ≠ wake | combined | delayed test | competition (rule-based) accuracy | $-0.82$ | $124$ | $.413$ | $-0.15$ |
| overall ≠ chance performance | combined | overall | competition (rule-based) accuracy | $-12.09$ | $182$ | $< .001^{***}$ | $-0.89$ |

*Note*. ***: $p < .001$; **: $p < .01$.

**Supplementary Information 4: Group summaries of reinforcement learning (RL) model prediction accuracies**

***Table S4A.*** Learning (2nd half) model prediction accuracies group summary.

| group | study type | $M$ | $SE$ |
|---|---|---|---|
| control morning | main | 85.23 | 1.46 |
| control evening | main | 86.23 | 0.83 |
| sleep | main | 85.11 | 0.94 |
| wake | main | 86.22 | 0.94 |
| overall | main | 85.70 | 0.52 |
| sleep | fMRI | 84.29 | 0.98 |
| wake | fMRI | 84.72 | 1.67 |
| overall | fMRI | 84.50 | 0.93 |
| control morning | combined | 85.23 | 1.46 |
| control evening | combined | 86.23 | 0.83 |
| sleep | combined | 84.84 | 0.70 |
| wake | combined | 85.77 | 0.83 |
| overall | combined | 85.44 | 0.45 |

***Table S4B.*** Explicit model prediction accuracies group summary.

| **group** | **study type** | $M$ | $SE$ |
|---|---|---|---|
| control morning | main | 69.71 | 1.86 |
| control evening | main | 67.31 | 1.29 |
| sleep | main | 72.90 | 1.39 |
| wake | main | 71.73 | 1.16 |
| overall | main | 70.78 | 0.72 |
| sleep | fMRI | 67.14 | 1.90 |
| wake | fMRI | 65.53 | 1.57 |
| overall | fMRI | 66.38 | 1.24 |
| control morning | combined | 69.71 | 1.86 |
| control evening | combined | 67.31 | 1.29 |
| sleep | combined | 70.98 | 1.17 |
| wake | combined | 69.86 | 1.00 |
| overall | combined | 69.82 | 0.64 |

***Table S4C.*** Implicit model prediction accuracies group summary.

| group | study type | $M$ | $SE$ |
|---|---|---|---|
| control morning | main | 64.00 | 1.69 |
| control evening | main | 66.31 | 1.22 |
| sleep | main | 66.52 | 1.01 |
| wake | main | 64.16 | 1.17 |
| overall | main | 65.26 | 0.62 |
| sleep | fMRI | 67.57 | 1.17 |
| wake | fMRI | 63.58 | 1.76 |
| overall | fMRI | 65.67 | 1.07 |
| control morning | combined | 64.00 | 1.69 |
| control evening | combined | 66.31 | 1.22 |
| sleep | combined | 66.87 | 0.77 |
| wake | combined | 63.98 | 0.97 |
| overall | combined | 65.35 | 0.54 |

***Table S4D.*** Categorization model prediction accuracies group summary.

| group | study type | $M$ | $SE$ |
|---|---|---|---|
| control morning | main | 81.29 | 2.02 |
| control evening | main | 83.59 | 1.62 |
| sleep | main | 81.76 | 1.14 |
| wake | main | 78.50 | 1.27 |
| overall | main | 81.03 | 0.74 |
| sleep | fMRI | 78.29 | 1.58 |
| wake | fMRI | 77.05 | 2.01 |
| overall | fMRI | 77.70 | 1.25 |
| control morning | combined | 81.29 | 2.02 |
| control evening | combined | 83.59 | 1.62 |
| sleep | combined | 80.60 | 0.94 |
| wake | combined | 78.06 | 1.07 |
| overall | combined | 80.31 | 0.64 |

**Supplementary Information 5: Group comparisons of reinforcement learning (RL) model prediction accuracies**

***Table S5A.*** Learning (2nd half) model prediction accuracies group comparisons.

| group (comparison) | study type | test type | outcome | $t$ | $df$ | $p$ | $d$ |
|---|---|---|---|---|---|---|---|
| control morning ≠ control evening | main | immediate test | accuracy learning model prediction | $-0.60$ | $55$ | $.549$ | $-0.16$ |
| sleep ≠ wake | main | delayed test | accuracy learning model prediction | $-0.83$ | $84$ | $.409$ | $-0.18$ |
| sleep ≠ control evening | main | evening training | accuracy learning model prediction | $-0.85$ | $69$ | $.401$ | $-0.20$ |
| wake ≠ control morning | main | morning training | accuracy learning model prediction | $0.60$ | $70$ | $.553$ | $0.14$ |
| overall ≠ chance performance | main | overall | accuracy learning model prediction | $69.23$ | $142$ | $< .001^{***}$ | $5.79$ |
| sleep ≠ wake | fMRI | delayed test | accuracy learning model prediction | $-0.23$ | $38$ | $.819$ | $-0.07$ |
| overall ≠ chance performance | fMRI | overall | accuracy learning model prediction | $37.07$ | $39$ | $< .001^{***}$ | $5.86$ |
| control morning ≠ control evening | combined | immediate test | accuracy learning model prediction | $-0.60$ | $55$ | $.549$ | $-0.16$ |
| sleep ≠ wake | combined | delayed test | accuracy learning model prediction | $-0.86$ | $124$ | $.393$ | $-0.15$ |
| sleep ≠ control evening | combined | evening training | accuracy learning model prediction | $-1.18$ | $90$ | $.240$ | $-0.27$ |
| wake ≠ control morning | combined | morning training | accuracy learning model prediction | $0.34$ | $89$ | $.733$ | $0.08$ |
| overall ≠ chance performance | combined | overall | accuracy learning model prediction | $78.44$ | $182$ | $< .001^{***}$ | $5.80$ |

*Note*. ***: $p < .001$.

***Table S5B.*** Explicit model prediction accuracies group comparisons.

| comparison | study type | test type | outcome | $t$ | $df$ | $p$ | $d$ |
|---|---|---|---|---|---|---|---|
| control morning ≠ control evening | main | immediate test | accuracy explicit model prediction | 1.07 | 55 | .290 | 0.28 |
| sleep ≠ wake | main | delayed test | accuracy explicit model prediction | 0.65 | 84 | .516 | 0.14 |
| overall ≠ chance performance | main | overall | accuracy explicit model prediction | 98.72 | 142 | < .001*** | 8.26 |
| sleep ≠ wake | fMRI | delayed test | accuracy explicit model prediction | 0.65 | 38 | .521 | 0.21 |
| overall ≠ chance performance | fMRI | overall | accuracy explicit model prediction | 53.70 | 39 | < .001*** | 8.49 |
| control morning ≠ control evening | combined | immediate test | accuracy explicit model prediction | 1.07 | 55 | .290 | 0.28 |
| sleep ≠ wake | combined | delayed test | accuracy explicit model prediction | 0.73 | 124 | .464 | 0.13 |
| overall ≠ chance performance | combined | overall | accuracy explicit model prediction | 109.95 | 182 | < .001*** | 8.13 |

*Note*. ***: $p < .001$.

***Table S5C.*** Implicit model prediction accuracies group comparisons.

| comparison | study type | test type | outcome | $t$ | $df$ | $p$ | $d$ |
|---|---|---|---|---|---|---|---|
| control morning ≠ control evening | main | immediate test | accuracy implicit model prediction | $-1.12$ | $55$ | $.269$ | $-0.30$ |
| sleep ≠ wake | main | delayed test | accuracy implicit model prediction | $1.52$ | $84$ | $.131$ | $0.33$ |
| overall ≠ chance performance | main | overall | accuracy implicit model prediction | $24.47$ | $142$ | $< .001$*** | $2.05$ |
| sleep ≠ wake | fMRI | delayed test | accuracy implicit model prediction | $1.93$ | $38$ | $.062$ | $0.61$ |
| overall ≠ chance performance | fMRI | overall | accuracy implicit model prediction | $14.65$ | $39$ | $< .001$*** | $2.32$ |
| control morning ≠ control evening | combined | immediate test | accuracy implicit model prediction | $-1.12$ | $55$ | $.269$ | $-0.30$ |
| sleep ≠ wake | combined | delayed test | accuracy implicit model prediction | $2.33$ | $124$ | $.021$* | $0.42$ |
| overall ≠ chance performance | combined | overall | accuracy implicit model prediction | $28.46$ | $182$ | $< .001$*** | $2.10$ |

*Note*. ***: $p < .001$; *: $p < .05$.

***Table S5D.*** Categorization model prediction accuracies group comparisons.

| comparison | study type | test type | outcome | $t$ | $df$ | $p$ | $d$ |
|---|---|---|---|---|---|---|---|
| control morning ≠ control evening | main | immediate test | accuracy categorization model prediction | $-0.89$ | $55$ | $.377$ | $-0.24$ |
| sleep ≠ wake | main | delayed test | accuracy categorization model prediction | $1.90$ | $84$ | $.060$ | $0.41$ |
| overall ≠ chance performance | main | overall | accuracy categorization model prediction | $42.14$ | $142$ | $< .001$*** | $3.52$ |
| sleep ≠ wake | fMRI | delayed test | accuracy categorization model prediction | $0.49$ | $38$ | $.630$ | $0.15$ |
| overall ≠ chance performance | fMRI | overall | accuracy categorization model prediction | $22.09$ | $39$ | $< .001$*** | $3.49$ |
| control morning ≠ control evening | combined | immediate test | accuracy categorization model prediction | $-0.89$ | $55$ | $.377$ | $-0.24$ |
| sleep ≠ wake | combined | delayed test | accuracy categorization model prediction | $1.78$ | $124$ | $.077$ | $0.32$ |
| overall ≠ chance performance | combined | overall | accuracy categorization model prediction | $47.06$ | $182$ | $< .001$*** | $3.48$ |

*Note*. ***: $p < .001$.

**Supplementary Information 6: Group summaries of feature selection reinforcement learning (RL) model prediction accuracies**

***Table S6A.*** Implicit feature selection model prediction accuracies group summary for the three features relevant for classification.

| group | type | $M$ | $SE$ |
|---|---|---|---|
| control morning | behavioral | 63.93 | 1.67 |
| control evening | behavioral | 66.28 | 1.21 |
| sleep | behavioral | 66.48 | 1.00 |
| wake | behavioral | 64.14 | 1.18 |
| overall | behavioral | 65.22 | 0.62 |
| sleep | fMRI | 67.71 | 1.19 |
| wake | fMRI | 63.37 | 1.72 |
| overall | fMRI | 65.65 | 1.07 |
| control morning | combined | 63.93 | 1.67 |
| control evening | combined | 66.28 | 1.21 |
| sleep | combined | 66.89 | 0.77 |
| wake | combined | 63.90 | 0.97 |
| overall | combined | 65.31 | 0.54 |

***Table S6B.*** Implicit feature selection model prediction accuracy differences group summary for the combined three features relevant for classification minus the average of the single features relevant for classification.

| group | type | $M$ | $SE$ |
|---|---|---|---|
| control morning | behavioral | 6.07 | 0.99 |
| control evening | behavioral | 5.40 | 0.69 |
| sleep | behavioral | 6.48 | 0.63 |
| wake | behavioral | 6.21 | 0.65 |
| overall | behavioral | 6.10 | 0.36 |
| sleep | fMRI | 7.81 | 0.62 |
| wake | fMRI | 5.19 | 0.85 |
| overall | fMRI | 6.57 | 0.55 |
| control morning | combined | 6.07 | 0.99 |
| control evening | combined | 5.40 | 0.69 |
| sleep | combined | 6.93 | 0.47 |
| wake | combined | 5.90 | 0.52 |
| overall | combined | 6.20 | 0.31 |

**Supplementary Information 7: Group comparisons of feature selection reinforcement learning (RL) model prediction accuracies**

***Table S7A.*** Implicit feature selection model prediction accuracies group comparisons for the three features relevant for classification.

| comparison | study type | test type | outcome | $t$ | $df$ | $p$ | $d$ |
|---|---|---|---|---|---|---|---|
| control morning ≠ control evening | behavioral | immediate test | accuracy three features implicit prediction | $-1.14$ | $55$ | $.258$ | $-0.30$ |
| sleep ≠ wake | behavioral | delayed test | accuracy three features implicit prediction | $1.51$ | $84$ | $.136$ | $0.32$ |
| overall ≠ chance performance | behavioral | overall | accuracy three features implicit prediction | $24.46$ | $142$ | $< .001$*** | $2.05$ |
| sleep ≠ wake | fMRI | delayed test | accuracy three features implicit prediction | $2.11$ | $38$ | $.041$* | $0.67$ |
| overall ≠ chance performance | fMRI | overall | accuracy three features implicit prediction | $14.61$ | $39$ | $< .001$*** | $2.31$ |
| control morning ≠ control evening | combined | immediate test | accuracy three features implicit prediction | $-1.14$ | $55$ | $.258$ | $-0.30$ |
| sleep ≠ wake | combined | delayed test | accuracy three features implicit prediction | $2.41$ | $124$ | $.017$* | $0.43$ |
| overall ≠ chance performance | combined | overall | accuracy three features implicit prediction | $28.43$ | $182$ | $< .001$*** | $2.10$ |

*Note*. ***: $p < .001$; *: $p < .05$.

***Table S7B.*** Implicit feature selection model prediction accuracy differences group comparisons for the combined three features relevant for classification minus the average of the single features relevant for classification.

| comparison | study type | test type | outcome | $t$ | $df$ | $p$ | $d$ |
|---|---|---|---|---|---|---|---|
| control morning ≠ control evening | behavioral | immediate test | accuracy three minus one features implicit prediction | $0.56$ | $55$ | $.581$ | $0.15$ |
| sleep ≠ wake | behavioral | delayed test | accuracy three minus one features implicit prediction | $0.30$ | $84$ | $.765$ | $0.06$ |
| overall ≠ chance performance | behavioral | overall | accuracy three minus one features implicit prediction | $-121.61$ | $142$ | $< .001$*** | $-10.17$ |
| sleep ≠ wake | fMRI | delayed test | accuracy three minus one features implicit prediction | $2.52$ | $38$ | $.016$* | $0.80$ |
| overall ≠ chance performance | fMRI | overall | accuracy three minus one features implicit prediction | $-78.42$ | $39$ | $< .001$*** | $-12.40$ |
| control morning ≠ control evening | combined | immediate test | accuracy three minus one features implicit prediction | $0.56$ | $55$ | $.581$ | $0.15$ |
| sleep ≠ wake | combined | delayed test | accuracy three minus one features implicit prediction | $1.45$ | $124$ | $.150$ | $0.26$ |
| overall ≠ chance performance | combined | overall | accuracy three minus one features implicit prediction | $-142.84$ | $182$ | $< .001$*** | $-10.56$ |

*Note*. ***: $p < .001$; *: $p < .05$.

**Supplementary Information 8: Group comparison (night-sleep > day-wake) of brain clusters activated during memory tests (correct > incorrect responses)**

***Table S8A.*** Group comparison (night-sleep > day-wake) of brain clusters activated during explicit memory test (correct > incorrect contrast).

| | | | MNI Coordinates | | |
|---|---|---|---|---|---|
| **Region Label** | **Extent** | ***t*** | **x** | **y** | **z** |
| R Precuneus (white matter) / R Middle Temporal Gyrus (white matter)* | 118 | 4.028 | 30 | −51 | 24 |
| L Middle Occipital Gyrus (white matter) / L Superior Occipital Gyrus (white matter) / L Superior Parietal Lobule (white matter)* | 236 | 3.630 | −25 | −55 | 32 |
| R Anterior Cingulate Cortex* | 42 | 3.590 | 18 | 24 | 30 |
| L Superior Frontal Gyrus (white matter) / L Precentral Gyrus (white matter)* | 31 | 3.441 | −23 | −19 | 54 |
| L Inferior Frontal Gyrus (p. Orbitalis) | 49 | 3.371 | −39 | 46 | −11 |
| L Caudate Nucleus (white matter)* | 145 | 3.224 | −27 | −7 | 28 |
| L Superior Medial Gyrus | 60 | 3.157 | −9 | 24 | 54 |
| R Olfactory cortex* | 85 | 3.147 | 2 | 8 | 4 |
| L Middle Frontal Gyrus (white matter) / L Inferior Frontal Gyrus (p. Triangularis) (white matter)* | 42 | 3.137 | −27 | 22 | 22 |
| L Superior Orbital Gyrus | 16 | 3.097 | −17 | 26 | −11 |
| R Superior Frontal Gyrus | 21 | 3.061 | 24 | 50 | 6 |
| L Superior Temporal Gyrus (white matter) / L Heschl's Gyrus (white matter)* | 10 | 3.005 | −33 | −37 | 14 |
| R Parahippocampal Gyrus | 21 | 2.958 | 16 | −9 | −19 |
| L Postcentral Gyrus (white matter) / L | 11 | 2.941 | −33 | −7 | 38 |

| | | | | | |
|---|---|---|---|---|---|
| Precentral Gyrus (white matter)* | | | | | |
| White matter* | 12 | 2.911 | −29 | −25 | 34 |
| L Middle Temporal Gyrus | 23 | 2.839 | −51 | −37 | −9 |
| R Middle Orbital Gyrus (white matter) / R Inferior Frontal Gyrus (p. Orbitalis) (white matter)* | 6 | 2.824 | 44 | 42 | −19 |

*Note*. Whole-brain MNI space group-level analyses. All clusters exhibited significant peak-level effects at $p_{\text{uncorr}} < .005$ and exceeded 5 voxels. Coordinates were labeled using `bspmview`. * indicates that the region label was selected manually as the cluster peak location was not within the voxels labeled by the atlas.

***Table S8B.*** Group comparison (night-sleep > day-wake) of brain clusters activated during implicit memory test (correct > incorrect contrast).

| | | | MNI Coordinates | | |
|---|---|---|---|---|---|
| **Region Label** | **Extent** | $t$ | **x** | **y** | **z** |
| L Cerebellum (IX)* | 11 | 3.172 | −13 | −43 | −45 |
| L Mid Cingulate Cortex (white matter)* | 6 | 2.845 | −17 | −13 | 54 |
| R Hippocampus | 10 | 2.801 | 34 | −17 | −13 |
| R Temporal Pole | 8 | 2.789 | 44 | 6 | −21 |
| L Parahippocampal Gyrus (white matter) / L Hippocampus (white matter)* | 9 | 2.749 | −15 | −21 | −9 |
| R Postcentral Gyrus* | 6 | 2.716 | 68 | −7 | 34 |

*Note*. Whole-brain MNI space group-level analyses. All clusters exhibited significant peak-level effects at $p_{\text{uncorr}} < .005$ and exceeded 5 voxels. Coordinates were labeled using `bspmview`. * indicates that the region label was selected manually as the cluster peak location was not within the voxels labeled by the atlas.

***Table S8C.*** Group comparison (night-sleep > day-wake) of brain clusters activated during categorization memory test (correct > incorrect contrast).

| | | | MNI Coordinates | | |
|---|---|---|---|---|---|
| **Region Label** | **Extent** | ***t*** | **x** | **y** | **z** |
| R Anterior Cingulate Cortex | 8 | 2.907 | 8 | 34 | 2 |
| R Putamen (white matter)* | 8 | 2.796 | 30 | −29 | 10 |

*Note*. Whole-brain MNI space group-level analyses. All clusters exhibited significant peak-level effects at $p_{\text{uncorr}} < .005$ and exceeded 5 voxels. Coordinates were labeled using `bspmview`. * indicates that the region label was selected manually as the cluster peak location was not within the voxels labeled by the atlas.

**Supplementary Information 9: Group comparison (night-sleep > day-wake) of brain clusters modulated by exemplar value representation difference during implicit memory test ($Q$-value modulation)**

***Table S9.*** Group comparison (night-sleep > day-wake) of brain clusters modulated by exemplar value representation difference during implicit memory test ($Q$-value modulation).

| | | | MNI Coordinates | | |
|---|---|---|---|---|---|
| **Region Label** | **Extent** | $t$ | **x** | **y** | **z** |
| R Cerebellum (IV-V) | 44 | 3.375 | 20 | −41 | −15 |
| L Anterior Cingulate Cortex (white matter)* | 28 | 3.221 | −17 | 16 | 36 |
| R Superior Orbital Gyrus | 18 | 3.182 | 16 | 44 | −7 |
| R Superior Temporal Gyrus | 49 | 3.151 | 46 | −11 | −9 |
| L Superior Orbital Gyrus | 7 | 3.103 | −19 | 48 | −9 |
| R Cerebellum (VIII) | 9 | 2.911 | 30 | −63 | −43 |

*Note*. Whole-brain MNI space group-level analyses. All clusters exhibited significant peak-level effects at $p_{\mathrm{uncorr}} < .005$ and exceeded 5 voxels. Coordinates were labeled using `bspmview`. * indicates that the region label was selected manually as the cluster peak location was not within the voxels labeled by the atlas.

### Supplementary Information 10: Group comparison (night-sleep > day-wake) of brain clusters activated during memory tests (correct > incorrect responses) and by exemplar value representation difference during implicit memory test (Q-value modulation)

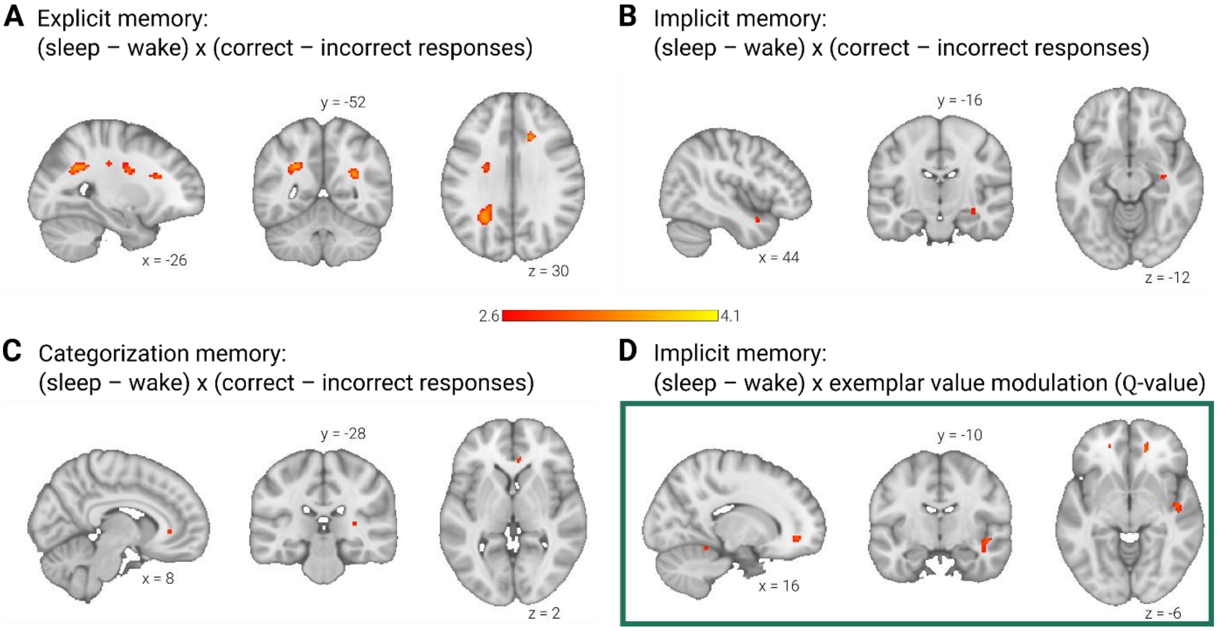


***Figure S10. Group comparison (night-sleep > day-wake) of neural activity during explicit, implicit, and categorization memory tests (correct > incorrect contrasts) and of neural activity modulated by exemplar value representation difference during implicit memory test (Q-value modulation).*** (A) Explicit memory test (sleep > wake) x (correct > incorrect responses): superior parietal lobule, precuneus, parahippocampal gyrus, frontal cortex and caudate nucleus (Supplementary Table S8A). (B) Implicit memory (sleep > wake) x (correct > incorrect responses): bilateral temporal lobe, including hippocampus (Supplementary Table S8B). (C) Categorization memory (sleep > wake) x (correct > incorrect responses): putamen, medial frontal cortex (Supplementary Table S8C). (D) Implicit memory (sleep > wake) x exemplar value modulation ($Q$-value): medial frontal cortex (Supplementary Table S9). (A to D) Whole-brain MNI space group-level analyses. Neural activity was measured using functional magnetic resonance imaging (fMRI). All clusters exhibited significant peak-level effects at $p_{\text{uncorr}} < .005$ and exceeded 5 voxels. No masking. Images are displayed in anatomical orientation.

**Supplementary Information 11: Explicit insight into the exemplar value rule**

***Table S11A.*** Hierarchy and combination rule rater agreement.

| rule | rater agreement (%) |
|---|---|
| hierarchy | 88 |
| combination | 96 |

***Table S11B.*** Relationship between implicit memory performance and insight into exemplar value rule summary.

| study type | rule | rule reported | $M_{res}$ | $SE_{res}$ |
|---|---|---|---|---|
| main | hierarchy | not reported | $-0.02$ | $0.01$ |
| main | hierarchy | reported | $0.02$ | $0.01$ |
| main | combination | not reported | $-0.02$ | $0.01$ |
| main | combination | reported | $0.02$ | $0.01$ |
| fMRI | hierarchy | not reported | $-0.01$ | $0.02$ |
| fMRI | hierarchy | reported | $0.01$ | $0.01$ |
| fMRI | combination | not reported | $-0.03$ | $0.02$ |
| fMRI | combination | reported | $0.02$ | $0.01$ |
| combined | hierarchy | not reported | $-0.02$ | $0.01$ |
| combined | hierarchy | reported | $0.01$ | $0.01$ |
| combined | combination | not reported | $-0.02$ | $0.01$ |
| combined | combination | reported | $0.02$ | $0.01$ |

***Table S11C.*** Relationship between implicit memory performance and insight into exemplar value rule comparison.

| comparison | study type | rule | outcome | $t$ | $df$ | $p$ | $d$ |
|---|---|---|---|---|---|---|---|
| rule not reported ≠ rule reported | main | hierarchy | implicit accuracy | $-2.68$ | $141$ | $.008$** | $-0.45$ |
| rule not reported ≠ rule reported | main | combination | implicit accuracy | $-2.92$ | $141$ | $.004$** | $-0.49$ |
| rule not reported ≠ rule reported | fMRI | hierarchy | implicit accuracy | $-0.85$ | $38$ | $.401$ | $-0.27$ |
| rule not reported ≠ rule reported | fMRI | combination | implicit accuracy | $-2.14$ | $38$ | $.039$* | $-0.69$ |
| rule not reported ≠ rule reported | combined | hierarchy | implicit accuracy | $-2.84$ | $181$ | $.005$** | $-0.42$ |
| rule not reported ≠ rule reported | combined | combination | implicit accuracy | $-3.61$ | $181$ | $< .001$*** | $-0.54$ |

*Note*. ***: $p < .001$; **: $p < .01$; *: $p < .05$.

***Table S11D.*** Participants' reported hierarchy and combination rules group summary.

| group | study type | hierarchy (%) | combination (%) |
|---|---|---|---|
| control morning | main | 39 | 25 |
| control evening | main | 48 | 38 |
| sleep | main | 55 | 57 |
| wake | main | 50 | 43 |
| sleep | fMRI | 52 | 62 |
| wake | fMRI | 63 | 53 |
| control morning | combined | 39 | 25 |
| control evening | combined | 48 | 38 |
| sleep | combined | 54 | 59 |
| wake | combined | 54 | 46 |

***Table S11E.*** Participants' reported hierarchy rules group comparisons.

| comparison | study type | test type | outcome | $X^2$ | *df* | *p* |
|---|---|---|---|---|---|---|
| control morning ≠ control evening | main | immediate test | hierarchy rule (count) | $0.17$ | $1$ | $.677$ |
| sleep ≠ wake | main | delayed test | hierarchy rule (count) | $0.05$ | $1$ | $.821$ |
| sleep ≠ wake | fMRI | delayed test | hierarchy rule (count) | $0.14$ | $1$ | $.713$ |
| control morning ≠ control evening | combined | immediate test | hierarchy rule (count) | $0.17$ | $1$ | $.677$ |
| sleep ≠ wake | combined | delayed test | hierarchy rule (count) | $0.00$ | $1$ | $1.00$ |

***Table S11F.*** Participants' reported combination rules group comparisons.

| comparison | study type | test type | outcome | $X^2$ | $df$ | $p$ |
|---|---|---|---|---|---|---|
| control morning ≠ control evening | main | immediate test | combination rule (count) | $0.59$ | $1$ | $.444$ |
| sleep ≠ wake | main | delayed test | combination rule (count) | $1.16$ | $1$ | $.281$ |
| sleep ≠ wake | fMRI | delayed test | combination rule (count) | $0.07$ | $1$ | $.785$ |
| control morning ≠ control evening | combined | immediate test | combination rule (count) | $0.59$ | $1$ | $.444$ |
| sleep ≠ wake | combined | delayed test | combination rule (count) | $1.56$ | $1$ | $.212$ |

### Supplementary Information 12: Reinforcement learning (RL) feature weights at the end of learning

***Table S12A.*** Reinforcement learning (RL) feature weights at the end of learning feature summary.

| feature | $M$ | $SE$ |
|---|---|---|
| shape | 0.48 | 0.01 |
| symbol | 0.48 | 0.01 |
| fill | 0.48 | 0.01 |
| frame (irrelevant) | 0.00 | 0.00 |

***Table S12B.*** Reinforcement learning (RL) feature weights at the end of learning feature comparisons.

| **comparison** | **outcome** | $t$ | $df$ | $p$ | $d$ |
|---|---|---|---|---|---|
| shape ≠ symbol | feature weight model prediction | $-0.75$ | $182$ | $.452$ | $-0.06$ |
| shape ≠ fill | feature weight model prediction | $-0.42$ | $182$ | $.678$ | $-0.03$ |
| shape ≠ frame | feature weight model prediction | $44.16$ | $182$ | $< .001^{***}$ | $3.26$ |
| symbol ≠ fill | feature weight model prediction | $0.37$ | $182$ | $.711$ | $0.03$ |
| symbol ≠ frame | feature weight model prediction | $40.34$ | $182$ | $< .001^{***}$ | $2.98$ |
| fill ≠ frame | feature weight model prediction | $41.41$ | $182$ | $< .001^{***}$ | $3.06$ |

*Note*. ***: $p < .001$.
